# Supplemental deinoxanthin ameliorates bone marrow microenvironmental impairments and recovers functional damage of bone marrow-retained cells in total body irradiation-exposed mice

**DOI:** 10.64898/2026.05.27.728296

**Authors:** Shankar Rijal, Kihyun Kim, Govinda Bhattarai, Junhyeok Kim, Bitna Kim, Young-Mi Jeon, Sung-Ho Kook, Jeong-Chae Lee

## Abstract

Total body irradiation (TBI) can impair the bone marrow (BM) microenvironment and dysregulate the fates of BM-residing cells by overproducing reactive oxygen species (ROS) and inflammatory mediators. This study aims to investigate the potency and mechanism of *Deinococcus radiodurans*-derived deinoxanthin (DEIX) in mitigating TBI-mediated injuries in the BM microenvironment and BM-resident cells. C57BL/6 mice were divided into control, TBI, TBI+DEIX, and/or DEIX groups, in which the mice were exposed to sub-lethal TBI (5 Gy) or in combination with oral DEIX supplementation (25 mg/kg body weight). While the DEIX’s effect on BM and BM-resident cells was determined after five weeks of TBI, RNA sequence profiling on the mouse group-derived BM cells was performed after two weeks of TBI. Supplementation with DEIX protected mice against TBI-mediated decrease in bone mineral density of trabecular bones. Supplemental DEIX suppressed BM microenvironmental impairment and the induction of oxidative stress and senescence in BM cells of TBI-exposed mice. That suppression was orchestrated by the DEIX-induced restoration of TBI-stimulated disorders in osteogenic, osteoclastogenic, and adipogenic activation in the BM. Ex vivo assays using BM cells supported the notion that DEIX restores TBI-mediated defects in BM cell function, including colony formation, migration, and differentiation. RNA sequence profiling demonstrated DEIX’s potency to modulate the expression of genes that regulate cellular and systemic immune responses, cell proliferation and differentiation, and bone metabolism. Collectively, our results highlight the roles and associated mechanisms of DEIX in mitigating TBI-mediated microenvironmental impairment and in regulating BM-resident cells.

## Introduction

Total body irradiation (TBI) is a standard treatment for cancer patients and those requiring bone marrow (BM) transplantation. However, TBI-derived radiotherapy has adverse effects, such as internal organ injury, BM microenvironmental impairment, abnormal hematopoietic development, and stem cell dysfunction [1–3]. TBI-mediated impairments in BM and BM-retained stem cells can cause irrecoverable systemic damage. Considerable findings indicate that TBI-derived BM damage is triggered by increases in reactive oxygen species (ROS), inflammatory mediators, and osteoclastogenic molecules in BM-residing hematopoietic stem cells (HSCs) and mesenchymal stem cells (MSCs) [4–7]. Abnormal and prolonged accumulation of cellular ROS induces stem cell senescence and self-renewal defects in BM [7]. TBI-derived BM injury also impairs bone mass accrual and homeostatic maintenance, and those impairments are associated with the characteristic of bone to absorb radiation more sensitively than the surrounding soft tissues [8, 9].

Because TBI-induced adverse effects pose significant challenges for radiotherapy, studies have focused on developing bioactive compounds to prevent or reverse TBI-mediated impairments. Many studies suggest that naturally occurring phenolic compounds exhibit potent bioactivity in ameliorating or protecting against TBI-mediated oxidative damage [10–12]. The bioactivity of phenolic compounds stems from their ability to scavenge intracellular ROS [13], maintain the stemness of skeletal stem cells [14], and activate anti-aging and anti-inflammatory signaling pathways [15]. Carotenoids are also potent antioxidants that exhibit biological, medicinal, and pharmacological activities [16]. Interest in the use of deinoxanthin (DEIX; (2R)-2,1’-dihydroxy-3’,4’-didehydro-1’,2’-dihydro-β,ψ-caroten-4-one) as a radioprotective antioxidant is growing. DEIX is a unique xanthophyll carotenoid that was first characterized from *Deinococcus radiodurans* (*D. radiodurans*). DEIX showed stronger antioxidant potential in scavenging singlet oxygen and hydrogen peroxide than other xanthophyll carotenoids [17]. DEIX also had radioprotective ability against even high-dose irradiation [18]. The antioxidant and radioprotective properties of DEIX are believed to be associated with its unique structural features [18]. We previously found that the addition of DEIX directly removes cellular ROS and recovers the imbalance between osteoblast and osteoclast activities in hydrogen peroxide-exposed BM cells [19]. Supplemental DEIX protected mice against TBI-mediated damage to growth, organs, survival, and circulating blood cell production in a mouse model [19]. Supplemental DEIX also recovered the values of bone volume (BV, mm^3^) and BV percentage (BV/TV, %) in femoral bones of TBI-exposed mice. In addition, our recent findings highlighted the potential of DEIX to protect against inflammatory tissue destruction in experimental animal models of periodontitis [20]. Taken together, previous reports and our recent findings suggest that supplemental DEIX might prevent or attenuate TBI-mediated impairments in BM and BM-resident stem cells. However, the protective efficacy and the underlying mechanism of DEIX in TBI-induced oxidative BM damage remain unclear.

Here, we examined the radioprotective potency of long-term DEIX supplementation on TBI-mediated injuries in BM and BM-residing mesenchymal lineage cells, along with the associated mechanisms. To address this, we used mouse models exposed to sub-lethal TBI (5 Gy), with or without oral administration of DEIX, as described previously [19]. We evaluated bone mineral density (BMD, g/cm^3^) in trabecular bone of femurs, maintenance of the BM microenvironment, induction of senescence and oxidative stress in MSCs, and stem functionality of BM-stromal cells (BMSCs) and BM monocytes (BMMs) in the mouse groups after five weeks of TBI. To further understand the mechanisms underlying DEIX-mediated radioprotection in BM and BM-derived mesenchymal cells, we isolated whole BM cells from the mouse groups two weeks after TBI and performed RNA sequencing. Consequently, our current findings demonstrate DEIX’s radioprotective efficacy and possible mechanisms in the TBI-exposed mouse model, and suggest its clinical usefulness as a radioprotective antioxidant.

## Material and methods

### Chemicals and laboratory equipment

The procedures for producing and identifying DEIX from *D. radiodurans* have been described previously [19]. Antibodies specific to cyclooxygenase-2 (COX2; BS1076), nuclear factor erythroid 2-related factor 2 (Nrf2; BS1258), and runt-related transcription factor 2 (RUNX2; BS2831) were purchased from Bioworld Technology, Inc. (St. Louis Park, MN, USA). The receptor activator of the nuclear factor (NF)-κB ligand (RANKL; ALX-804-243) was purchased from Enzo Life Sciences, Inc. (Farmingdale, NY, USA). Osterix (ab209484), osteopontin (OPN; ab8448), and γ-H_2_AX (ab26350) antibodies were obtained from Abcam (Cambridge, UK). Antibodies specific to 8-hydroxy-2^’^-deoxyguanosine (8-OHdG; sc-66036), cathepsin K (sc-4835), peroxisome proliferator-activated receptor γ (PPARγ; sc-390740), and β-actin (sc-47778) were purchased from Santa Cruz Biotechnology (Santa Cruz, CA, USA). Osteocalcin (OCN; AB10911) antibody was purchased from Millipore Corporation (Temecula, CA, USA). Fetal bovine serum (FBS) and antibiotic-antimycotic (2441713) were purchased from HyClone Laboratories (Logan, UT, USA) and Gibco (Life Technologies, Carlsbad, CA, USA), respectively. Unless specified otherwise, other chemicals, antibodies, and laboratory consumables were purchased from Sigma-Aldrich Co. LLC (St. Louis, MI, USA), Abcam, and Falcon Labware (Becton-Dickinson Biosciences, Franklin Lakes, NJ, USA) or SPL Life Sciences (Pocheon, Republic of Korea), respectively.

### Animal and ethics statement

C57BL/6 mice (male, six weeks old) were purchased from Damul Science (Daejeon, Republic of Korea) and acclimatized for seven days before use. During the experimental periods, all animals were housed at 22 ± 1 °C, 55% ± 5% humidity, and a 12 h light/dark autocycle with *ad libitum* feeding in the Animal Center of the School of Dentistry, Jeonbuk National University (LML 18-620). We carried out this study in strict accordance with the recommendations in the Animal Care and Use Guide of Jeonbuk National University and ARRIVE guidelines 2.0 (https://arriveguidelines.org). The University Committee on Ethics approved all experimental procedures in the Care and Use of Laboratory Animals (NON2024-043-003 and NON2025-153-002).

### DEIX administration, TBI, and sample preparation

To evaluate the long-term supplemental effect of DEIX on the BM microenvironment and BM-residing mesenchymal cells, mice were randomly divided into three groups (*n* = 6/group): non-TBI mice (control group), TBI-exposed mice who received oral gavage of 100 μL of olive oil (vehicle) (TBI group), and TBI-exposed mice who were orally treated with DEIX (TBI+DEIX group). The TBI+DEIX group received DEIX (25 mg/kg body weight) once per day for 42 consecutive days, from seven days before to 35 days after TBI, and the TBI group received only the vehicle on the same days. The TBI and TBI+DEIX groups were exposed to 5 Gy TBI with γ-rays by regulating the dosage time (0.66 Gy/min) using the radioactive half-life of γ-rays on a rotating platform (Model IR-221, MDS Nordion, Ottawa, Canada). The amount of supplemental DEIX was adjusted as the mice’s body weights changed. The oral gavage of olive oil itself (100 μL) didn’t cause any side effects to mice [21]. Blood samples, long bones, and whole BM cells were collected 12 h after the final DEIX administration and used for in vivo and ex vivo analyses. To investigate the mechanism underlying DEIX-induced radioprotection, mice were divided into four groups (*n* = 3/group): control, TBI, TBI+DEIX, and DEIX. In these groups, whole BM cells were harvested at 14 days post-TBI from the femoral and tibial bones by flushing them with PBS using a 5 mL syringe after cutting the ends of the bones [21]. Each BM sample was treated with TRIzol reagent (Invitrogen) and stored at -20 °C until RNA sequence profiling. In the collection of samples, all groups of mice were euthanized with CO_2_ in a chamber, following the guidelines for the Euthanasia of Animals (2020), the American Veterinary Medical Association (AVMA).

### Histological analyses

The femoral bones were fixed in 4% paraformaldehyde for 48 h and then subjected to an additional decalcification in 10% EDTA at 4 °C for three weeks. The tissues were dehydrated in an alcohol series, embedded in paraffin, and sectioned at 5.0 µm. For hematoxylin and eosin (H & E) staining, tissue sections were treated with hematoxylin (Gill No. 3) before being counterstained with 0.25% Eosin Y stain (Thermo Fisher Scientific, Waltham, MA, USA). A portion of the femoral sections was subjected to tartrate-resistant acid phosphatase (TRAP) staining using a leukocyte acid phosphatase kit (Thermo Fisher Scientific), followed by counterstaining with hematoxylin. The expression patterns of adiponectin, cathepsin K, COX2, γ-H_2_AX, Nrf2, 8-OHdG, osterix, OCN, PPARγ, RUNX2, superoxide dismutase 1 (SOD-1), and heme oxygenase 1 (HO-1) in the tissue samples were evaluated by immunohistochemistry (IHC). In the IHC assay, tissue sections were stained with each primary antibody (1:200–500 dilutions) specific to the target molecules, and their expression patterns were determined using rabbit-anti- or mouse-anti-Vectastain ABC DAB-HRP kits (CAS:SK-4100, Vector Laboratories, Burlingame, CA, USA). All procedures for TRAP and IHC staining followed the relevant manufacturer’s instructions, and the stained sections were observed and photographed under a Motic slide scanner (Motic Sci & Tech Co., Ltd., Scottsdale, AZ, USA).

### Micro-computed tomography (μCT) analysis

The femurs from the mouse groups were scanned using a desktop scanner (1076 Skyscan Micro-CT; Bruker, Kontich, Belgium) and analyzed with CTAn software (ver. 1.18) as described elsewhere [21]. In brief, the X-ray source was set to 75 kV and 100 μA, with a pixel size of 18 μm. The image slices were reconstructed using a cone-beam reconstruction software (NRecon software ver. 1.6.10.5). Based on the reconstructed three-dimensional (3D) μCT images, BMD (g/cm^3^) was determined by converting the attenuation data for the volume of interest into Hounsfield units and BMD units using phantoms (Skyscan) with a standard density corresponding to mouse bone.

### Measurement of serum RANKL and osteoprotegerin (OPG) levels

The levels of RANKL and OPG in sera were measured by enzyme-linked immunosorbent assay (ELISA) using mouse-anti-RANKL (ab100749, Abcam) and -OPG ELISA kits (ab100733, Abcam) in a microplate reader (SPECTROstar® Nano, BMG LABTECH, Ortenberg, Germany). All procedures for this assay followed the manufacturer’s instructions.

### Flow cytometric analysis

Immediately after red blood cell removal, we analyzed mouse-derived BM cells by multicolor flow cytometry (BD Aria III, BD Biosciences) and identified cell populations phenotypically using FlowJo software (FLOWJO, Ashland, OR, USA). Unless specified otherwise, antibodies used in this analysis were purchased from BD Biosciences. We characterized and defined BM MSCs (CD29^+^CD105^+^LSK) using a PE-Cy7-conjugated lineage cocktail, APC-Cy7-conjugated anti-Sca-1 (eBioscience, Waltham, MA, USA), PE-conjugated anti-CD29, and APC-conjugated anti-CD105 antibodies. To analyze the levels of cellular ROS, the lineage marker-conjugated MSCs were stained with MitoSox^TM^ Red (Invitrogen, Carlsbad, CA, USA). A portion of these cells was also stained with 5-dodecanoylaminofluorescein di-β-D-galactopyranoside (C_12_FDG; Molecular Probes, Eugene, OR, USA) to evaluate the level of senescence-associated β-galactosidase (SA-β-gal) activity in relation to TBI exposure and/or DEIX supplementation. In addition, the level of p16^INK4a^ (Santa Cruz Biotechnology) in the BM MSCs was determined by flow cytometry using Alexa Fluor 488-conjugated antibodies.

### Western blot analysis

The mouse group-derived BM cells were lysed in a cocktail buffer containing protease/phosphatase inhibitor (Cell Signaling Technology, Danvers, MA, USA). The protein extracts (15–20 µg/sample) were separated through sodium dodecyl sulfate-polyacrylamide gel electrophoresis on 10–12% gels and electroblotted onto polyvinylidene difluoride membranes. The blots were washed with a buffer including 10 mM Tris-HCl (pH 7.6), 150 mM NaCl, and 0.05% Tween-20 and blocked in 5% skim milk for 1 h before incubation with primary antibody specific to PPARγ (1:500), RUNX2 (1:500), OPN (1:500), or β-actin (1:2500). The immunoreactive bands on the membranes were visualized using an enhanced peroxidase detection kit (ELPIS-Biotech, Daejeon, Republic Korea) and exposed to a chemiluminescence imaging system (Vilber Lourmat, Collegien, France).

### RNA isolation and real-time quantitative polymerase chain reaction (RT-qPCR) assay

In brief, total RNA was extracted from BM cells using TRIzol reagent (Invitrogen), and RNA samples (1 μg/sample) were used for cDNA synthesis with the AmpiGene cDNA Synthesis Kit (Enzo Life Sciences) according to the manufacturer’s instructions. RT-qPCR was performed with Power SYBR Green PCR Master Mix (Applied Biosystems, Waltham, MA, USA) and ABI StepOnePlus Real-Time PCR System (Applied Biosystems). The thermocycling conditions included a pre-denaturation at 95 °C for 10 min and amplification using three-step cycles of denaturation at 95 °C for 15 sec, annealing at 60 °C for 30 sec, and extension at 72 °C for 30 sec for 40 cycles. Oligonucleotide primers used in this study were designed as gagggactatggcgtcaaaca and ggatcccaaaagaagctttgc (XM_006523548.2) for *RUNX2* and acggacagctggcacaccag and ctcacacactcggttgtggg (NM_008764.3) for *OPN*. Glyceraldehyde-3-phosphate dehydrogenase (GAPDH) was used as the endogenous reference for quantification.

### Isolation and culture of BM cells for ex vivo assay

The mouse group-derived BM cells were resuspended in α-minimum essential medium (αMEM, Thermo Fisher Scientific) and centrifuged at 2,000×*g* for 3 min. The pellets were spread onto 60-mm culture plates and incubated in growth medium (αMEM supplemented with 2 mM glutamine, 100 IU/mL penicillin G, 100 μg/mL streptomycin, and 20% FBS). On the second day, the non-adherent supernatant cells were collected and used as BMMs, whereas the adherent cells were incubated further and used as BMSCs.

### Assays for ex vivo colony formation, migration, and differentiation of BMSCs

The effect of supplemental DEIX on the functioning of BMSCs derived from the mouse groups was examined by evaluating their ex vivo abilities to form colonies, migrate, and differentiate into osteocytes or adipocytes. Briefly, BMSCs were seeded onto 6-well culture plates (1 × 10^3^ cells/well) in growth medium. After 14 days of incubation, adherent cells were fixed in 10% formalin for 10 min and stained with 0.5% crystal violet dissolved in 100% methanol. Colonies that had formed were photographed using a light microscope (EL-Einsatz 451888, Carl Zeiss, Ostalbkreis, Germany), and colonies containing more than 50 cells were counted. To evaluate the effect of DEIX on cell migration, mouse group-derived BMSCs (5 × 10^5^ cells/well) were spread onto 6-well culture dishes in growth medium. After 12 h of incubation, the culture medium was replaced with serum-free medium. The cells were incubated for an additional 4 h, and then the bottoms of the culture plates were scraped to create a 1-mm-wide wound area. After scratching, cultures were fed with growth medium containing 1% FBS.

After 24 h of incubation, the scraped areas were photographed using a light microscope (EL-Einsatz 451888). Healing of the gap was analyzed in five randomly chosen fields in each dish, and the results are represented as the area (%) of migrated cells. Alternatively, BMSCs (2 × 10^5^ cells/well) were put into 12-well culture plates and incubated in osteogenic medium supplemented with 5% FBS and DAG (100 nM **<u>d</u>**examethasone, 50 µM **<u>a</u>**scorbic acid, and 10 mM β-**<u>g</u>**lycerophosphate). The culture medium was replaced every three days during the incubation. After 21 days of incubation, the degree of mineralization was determined by staining the cells with alizarin red S (ARS) after 20 min of fixation in 70% ethanol. The red dye-stained cells were photographed using a light microscope (EL-Einsatz 451888). The ARS-stained cells were also treated with 10% acetylpyridinium chloride, and the amount of red dye was quantified by measuring the dye-specific optical density at 405 nm using a microplate reader (SPECTROstar® Nano). A portion of BMSCs was also seeded onto 12-well culture plates (2 × 10^5^ cells/well) and incubated in adipogenic medium (α-MEM supplied with 5% FBS, 500 nM dexamethasone, 500 µM 3-isobutyl-l-methylxanthine, 100 µM indomethacin, and 10 µg/mL insulin). After 14 days of incubation, the cells were rinsed twice with PBS, fixed in a 10% formaldehyde solution for 1 h, and stained with oil red O (ORO) solution (Sigma-Aldrich Co., LLC) for 15 min. Red lipid droplets were observed in ORO-stained cells using a light microscope (EL-Einsatz 451888). Cellular lipid accumulation in ORO-stained cells was also quantified by measuring the dye-specific absorbance at 490 nm using a microplate reader.

### Assay for ex vivo osteoclastogenic differentiation of BMMs

BMMs derived from the mouse groups were seeded into 12-well culture plates (1 × 10^4^ cells/well) in the presence of 30 ng/mL murine macrophage colony-stimulating factor (M-CSF) and 50 ng/mL RANKL. After 5 days of incubation, the cultures were fixed with 4% paraformaldehyde in PBS and stained with a TRAP staining kit (Cosmo Bio Co., Tokyo, Japan) according to the manufacturer’s instructions. The TRAP-stained cells were photographed using a light microscope, and multinucleated cells with more than three nuclei were counted as osteoclasts. The mean osteoclast diameter was determined from the optical images.

### mRNA sequencing and functional enrichment analysis

Total RNAs were isolated from mouse group-derived BM cells using TRIzol (Invitrogen), and RNA purity for all samples was checked using a NanoVue (GE Healthcare, Chicago, IL, USA). Each RNA sample had an A260-A280 ratio greater than 1.8 and an A260-A230 ratio greater than 2.0. The RNA sequence analysis was performed at LAS (Kimpo, Republic of Korea) following the procedures for processing, read mapping, expression quantification, differentially expressed gene (DEG) analysis, functional annotation, and visualization. For preprocessing and genome mapping, potential sequencing adapters and low-quality bases in the raw reads were trimmed using Skewer (ver 0.2.2). The cleaned, high-quality reads, after trimming low-quality bases and sequencing adapters, were mapped to the reference genome using STAR (version 2.5). The strand-specific library option, --library-type=fr-first strand, was applied in the mapping process. To quantify mapped reads on the reference genome into gene expression values, Cuffquant in Cufflinks (ver 2.2.1) with the strand-specific library option --library-type=fr-first-strand and other default options was used. The gene annotation of the reference genome mm10 from the UCSC Genome Browser (https://genome.ucsc.edu) in GTF format was used as the gene models. The expression values were calculated in Fragments Per Kilobase of transcript per Million fragments mapped (FPKM) units. The DEGs between the two selected biological conditions were analyzed using Cuffdiff in the Cufflinks package with the strand-specific library option --library-type=fr-first-strand and other default options. To compare the expression profiles among the samples, the normalized expression values of a few hundred selected DEGs were clustered using in-house R scripts. The scatter plots of gene expression values and the volcano plots of expression fold changes and *p*-values between the two selected samples were generated using in-house R scripts. To get the insights on the biological functional role of the differential gene expression between the compared biological conditions, a gene set overlapping test between the analyzed DEGs and functionally categorized genes, including biological processes of Gene Ontology (GO), Kyoto Encyclopedia of Genes and Genomes (KEGG) pathways, and other functional gene sets, was performed by g:Profiler2 (ver 0.2.0).

### Statistical analysis

All data are represented as the mean ± standard deviation. Differences between the two groups were analyzed by an unpaired Student’s *t*-test using the GraphPad Prism (ver. 9.5) program (Boston, MA, USA). A value of *p* < 0.05 was considered statistically significant.

## Results

### DEIX recovers microenvironmental defect and lipid accumulation in the trabecular bone of TBI-exposed mice

To better understand the inhibitory effect of supplemental DEIX on TBI-induced decreases in BV (mm^3^) and BV/TV (%) values in the femoral bone [19], we evaluated the BMD value in the trabecular bone of TBI-exposed mice. We compared it with that of TBI+DEIX and untreated control mice. Similar to previous findings [19], the 2D (Fig. 1A) and 3D μCT (Fig. 1B) analyses indicated lower bone mass in trabecular bone of the TBI group than in the control and TBI+DEIX groups five weeks post-TBI. When bone quality was evaluated from 3D images, the TBI group had significantly lower BMD (g/cm3; *p* < 0.0001) than the control group. This decrease was significantly (*p* < 0.001) restored in the TBI+DEIX group (Fig. 1C). There was also a significant difference (*p* < 0.05) in BMD value between the control and TBI+DEIX groups. TBI-mediated BMD loss and its inhibition by supplemental DEIX were supported by the H & E staining, in which the TBI group exhibited relatively less bone mass in both the epiphysis and metaphysis of trabecular bone than the control and DEIX+TBI groups (Fig. 2A). The TBI group also showed significantly less area (%) of trabecular bone (Fig. 2B) and a higher number of lipid droplets than the control and DEIX+TBI groups (Fig. 2C). Consistent with this, IHC results exhibited greater expression of adiponectin (Fig. 2D) and PPARγ (Fig. 2E) in the BM of TBI group compared with the control and TBI+DEIX groups. These results suggest that TBI-mediated impairments in bone mass accrual and the BM microenvironment are orchestrated by lipid accumulation and that these impairments are recovered by supplementation with DEIX.

**Fig. 1.**
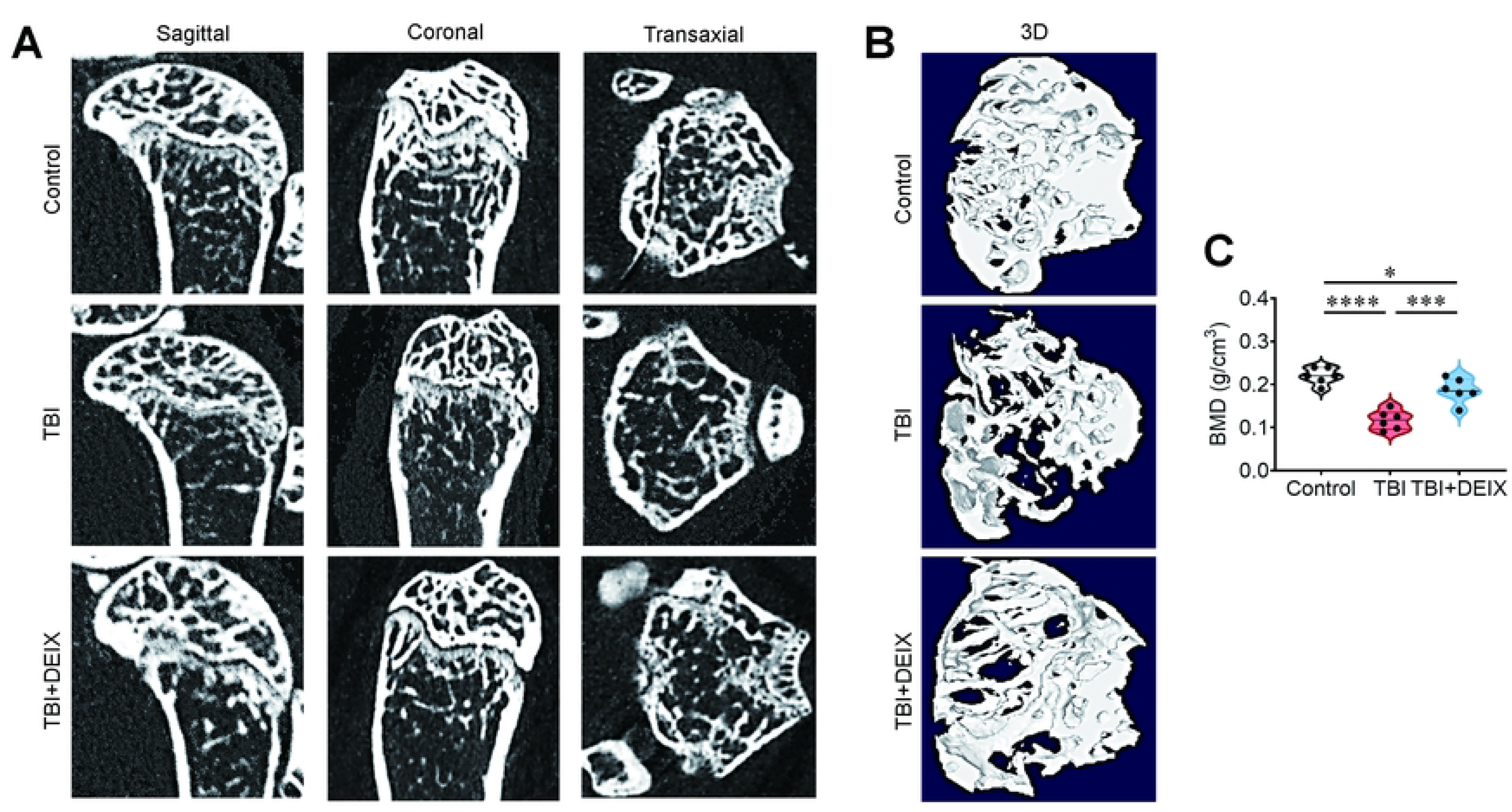
DEIX protects mice against TBI-mediated decrease in the BMD value of trabecular bone. (**A**) The 2D and (**B**) 3D μCT images showing trabecular bone of femurs isolated from the mouse groups five weeks after TBI. (**C**) The 3D image-based BMD (g/cm^3^) value in the trabecular region of femoral bones (*n* = 6). \**p* < 0.05, \*\*\**p* < 0.01, and \*\*\*\**p* < 0.0001 in an unpaired Student’s *t*-test.

**Fig. 2.**
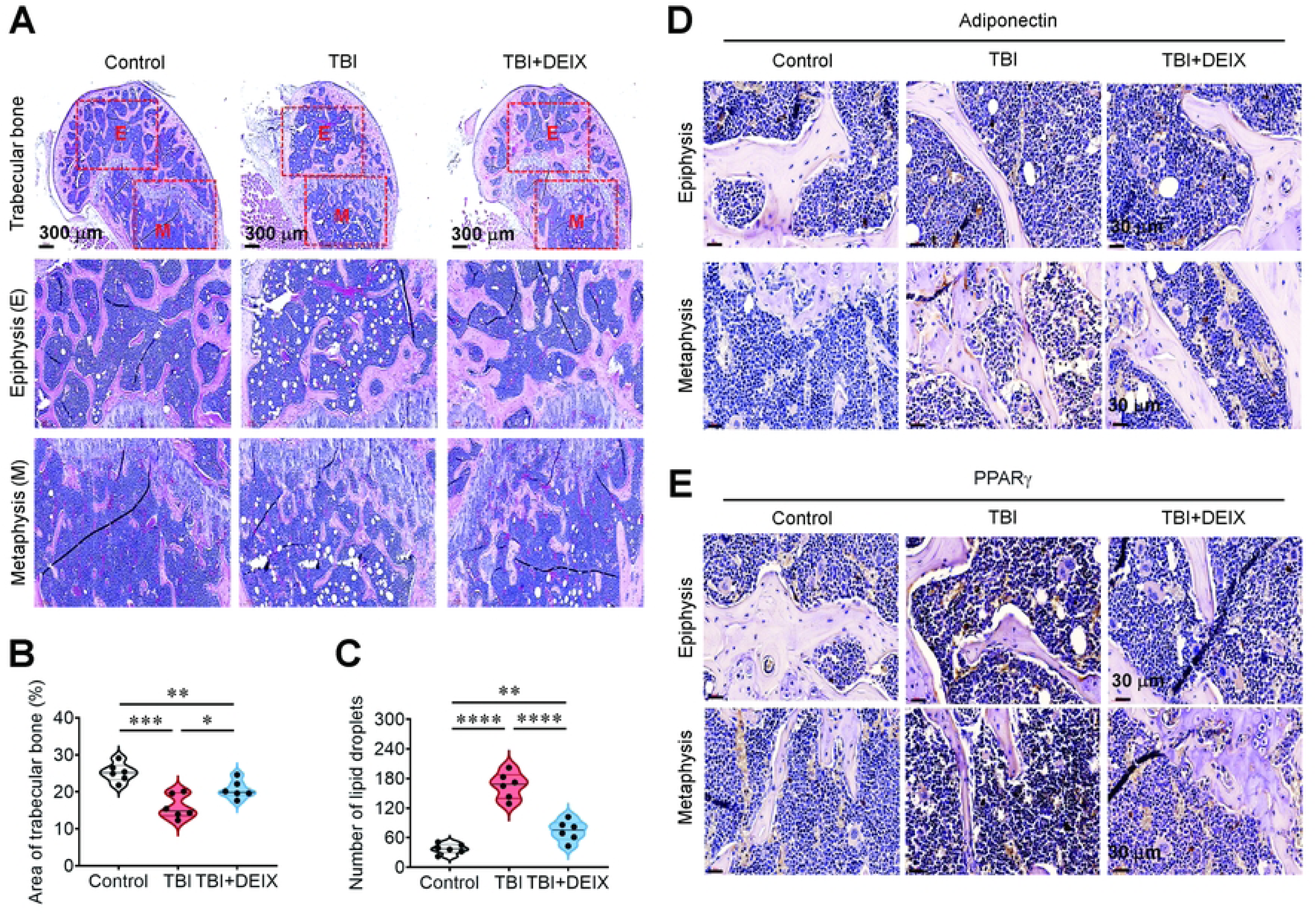
DEIX recovers bone mass loss and lipid accumulation in trabecular bones of TBI-exposed mice. (**A**) H & E staining images exhibiting bone mass accrual and lipid droplets in trabecular bone from the mouse groups five weeks post-TBI. (**B**) The area (%) of trabecular bone and (**C**) number of lipid droplets were determined from the H & E images (*n* = 6). Levels of (**D**) adiponectin and (**E**) PPARγ were determined in the BM of the mouse groups by IHC five weeks post-TBI. Representative data from six different samples are shown. \**p* < 0.05, \*\**p* < 0.01, \*\*\**p* < 0.001, and \*\*\*\**p* < 0.0001 in an unpaired Student’s *t*-test.

### Supplemental DEIX suppresses TBI-induced osteoclastic activation at local and systemic levels

Because an imbalanced osteogenic and osteoclastic activation is closely associated with TBI-mediated BM impairment and bone mass loss, we evaluated the effect of DEIX on the expression patterns of several osteogenic and osteoclastic molecules in the BM of mouse groups using IHC. The results indicated that BM of the TBI group exhibits the expression of RUNX2 (Fig. 3A) and osterix (Fig. 3B) similar to those of the control and TBI+DEIX groups. In contrast, lower OCN (Fig. 3C) and greater expression of cathepsin K (Fig. 3D) were found in the BM of the TBI group compared with the control and TBI+DEIX groups. When the serum levels of osteoclastogenesis-related proteins were determined, the TBI group had significantly higher RANKL levels than the control and TBI+DEIX groups (Fig. 3E), but not OPG (Fig. 3F). The TRAP-stained optical images also represented the TBI-stimulated osteoclast formation and its inhibition by supplemental DEIX (Fig. 3G). Furthermore, the TBI group showed significantly greater TRAP-positive areas (%) in both trabecular and cortical bone, compared with the control and TBI+DEIX groups (Fig. 3H). Our current findings indicate that, in addition to lipid accumulation, a skewed differentiation toward osteoclasts in the BM is associated with TBI-induced BM complications, and that the imbalance between osteoblast and osteoclast activities is restored by oral administration of DEIX.

**Fig. 3.**
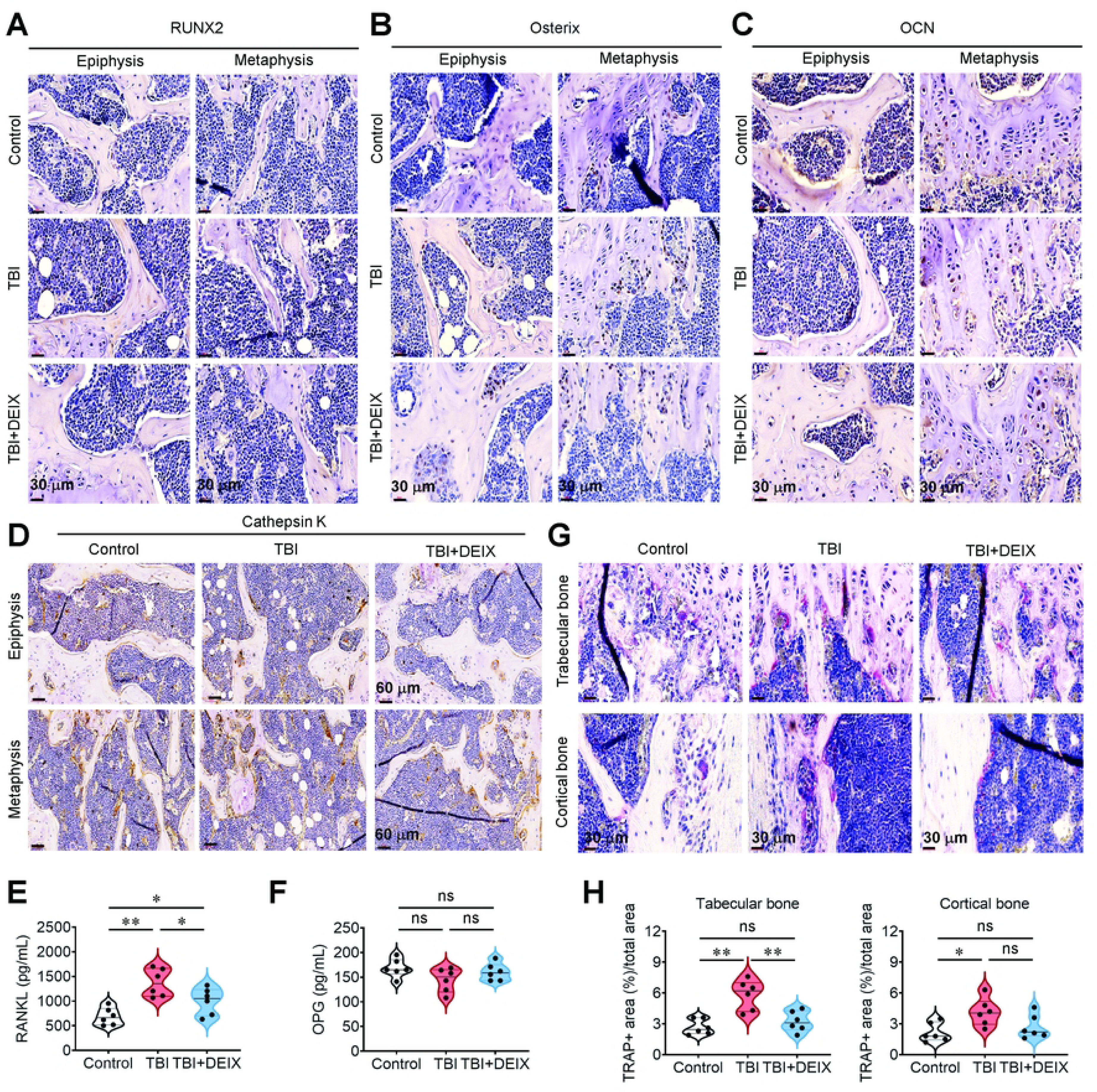
DEIX suppresses TBI-mediated skewed differentiation into osteoclasts in the BM of TBI-exposed mice. IHC results showing the levels of (**A**) RUNX2, (**B**) osterix, (**C**) OCN, and (**D**) cathepsin K in the BM of the mouse groups five weeks post-TBI. Representative data from six different mice are shown. Levels of (**E**) RANKL and (**F**) OPG in sera from the mouse groups were determined by ELISA five weeks after TBI (*n* = 6). (**G**) TRAP staining images showing osteoclast formation in trabecular and cortical bones from the mouse groups five weeks after TBI, along with (**H**) the area (%) positively stained by TRAP in trabecular and cortical bones (*n* = 6). \**p* < 0.05 and \*\**p* < 0.01 in an unpaired Student’s *t*-test. ns, not significant.

### DEIX inhibits the expression of DNA damage and inflammation-associated markers, but restores the levels of antioxidant molecules in the BM of TBI-exposed mice

IHC results revealed greater expression of 8-OHdg (Fig. 4A), γ-H_2_AX (Fig. 4B), HO-1 (Fig. 4C), and COX2 (Fig. 4D) in the BM of the TBI group compared with that of the control and TBI+DEIX groups. However, the BM of the TBI group showed lower expression of SOD-1 (Fig. 4E) and Nrf2 (Fig. 4F) than did the control and TBI+DEIX groups. These results indicate that TBI induces DNA damage, an inflammatory response, and decreased activity of the antioxidant defense system in the BM, and that this induction can be diminished or restored by DEIX supplementation.

**Fig. 4.**
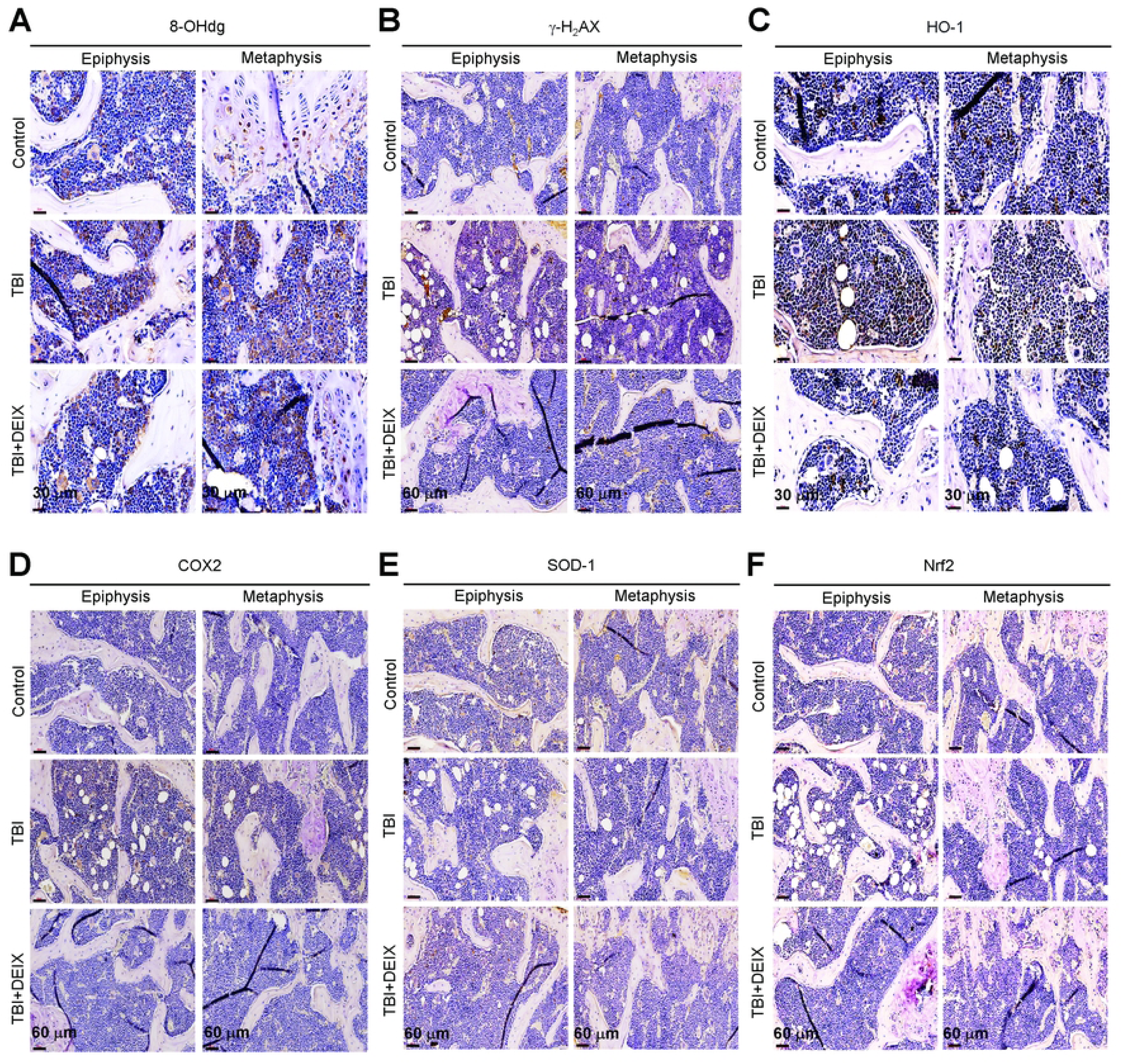
DEIX inhibits oxidative damage and inflammatory response and recovers the antioxidant defense system in the BM of TBI-exposed mice. IHC staining images exhibiting the expression patterns of (**A**) 8-OHdg, (**B**) γ-H_2_AX, (**C**) HO-1, (**D**) COX2, (**E**) SOD-1, and (**F**) Nrf2 in the BM of the control, TBI, and TBI+DEIX groups five weeks post-TBI.

### DEIX inhibits TBI-induced senescence and oxidative stress in BM MSCs and recovers TBI-derived functional loss of BMSCs and BMMs

Studies show that TBI impairs the function of BM-resident mesenchymal lineage cells primarily by inducing oxidative stress and senescence in a dose-dependent manner [22, 23]. Hence, we next explored whether the DEIX-derived beneficial effects are associated with its potency to inhibit senescence, oxidative damage, and/or functional defect of BM-residing mesenchymal cells. To this end, we determined the levels of BM MSCs that were positive for senescence (C_12_FDG and p16) or ROS-specific markers (MitoSox) by flow cytometry five weeks post-TBI. We also compared the capacities of BMSCs to form colonies, migrate, and differentiate into osteoblasts, adipocytes, and osteoclasts in relation to the TBI and/or DEIX supplementation. The total number of MSCs in the BM was not changed by TBI exposure, DEIX supplementation, or both in the mouse groups (Fig. 5A). However, the TBI group exhibited approximately 2-fold higher MSC levels positive to C_12_FDG (Fig. 5B), p16 (Fig. 5C), or MitoSox (Fig. 5D) when compared with the control and DEIX+TBI groups. The colony-forming activity of TBI group-derived BMSCs was significantly (*p* < 0.05) lower than that in control-or TBI+DEIX group-derived BMSCs (Fig. 5E). The TBI mice-derived BMSCs also showed a lower ability to migrate than the cells derived from the control or TBI+DEIX group (Fig. 5F). In addition, the TBI mice-derived BMSCs had a lower potency to differentiate into mineralized cells than the cells derived from the control or TBI+DEIX group (Fig. 6A). Similar to ARS staining results, the TBI group-derived BMSCs showed lower expression of RUNX2 and OPN at protein (Fig. 6B) and mRNA levels (Fig. 6C) compared with the control or TBI+DEIX group-derived cells. However, BMSCs derived from the TBI group showed greater potential to differentiate into adipocytes (Fig. 6D) and form lipid droplets than those derived from the control or TBI+DEIX group (Fig. 6E). TBI-stimulated lipid accumulation in BMSCs and its suppression by DEIX were supported by measuring the ORO dye-specific optical density (Fig. 6F), as well as by determining the protein level of PPARγ (Fig. 6G). The TBI group-derived BMMs exhibited greater ability to differentiate into multinucleated osteoclasts than those derived from the control or TBI+DEIX group (Fig. 6H). The TBI-stimulated osteoclast formation and its suppression by DEIX were supported by determining the number (Fig. 6I) and diameter of the osteoclasts that formed (Fig. 6J). These results indicate that TBI-mediated defects in the fate and function of BM-resident mesenchymal lineage cells are accompanied by oxidative damage and senescence induction, and that these defects are reversible with DEIX supplementation.

**Fig. 5.**
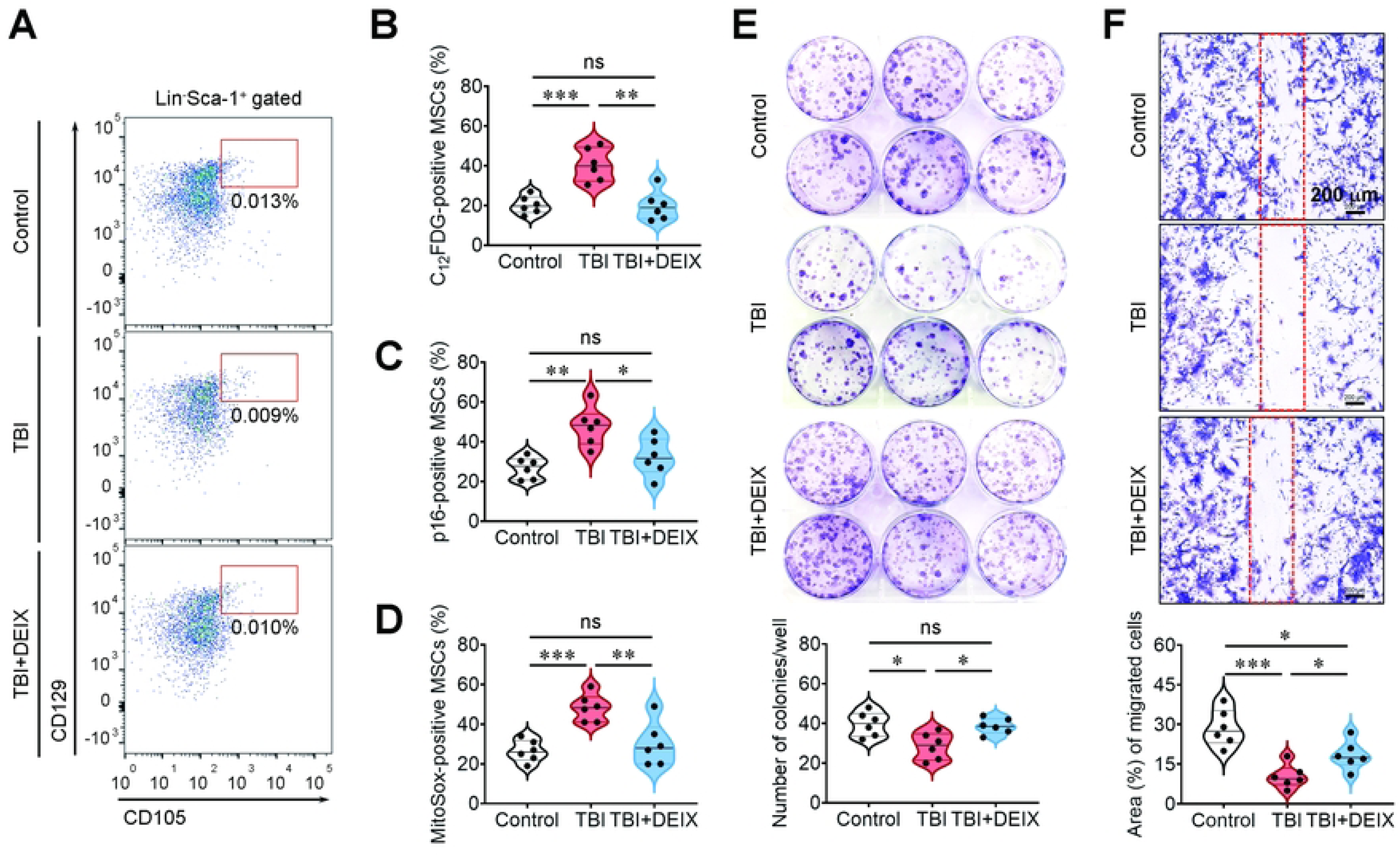
DEIX recovers TBI-induced senescence and oxidative stress in BM MSCs and restores colony-forming and migrating potencies of the mouse group-isolated BMSCs. Flow cytometric analysis was performed on BM-resident MSCs five weeks post-TBI. (**A)** The percentage of MSCs and the mean numbers (%) of MSCs positive for (**B)** C_12_FDG, (**C)** p16, or (**D)** MitoSox are shown (*n* = 6). BMSCs were isolated from the mouse groups five weeks post-TBI, and their colony-forming and migratory abilities were analyzed. (**E**) Photographs showing colony formation of BMSCs derived from the mouse groups, along with the number of colonies per well (*n* = 6). (**F**) Migration pattern of BMSCs and the area (%) of migrated BMSCs in the mouse groups (*n* = 6). \**p* < 0.05, \*\**p* < 0.01, and \*\*\**p* < 0.001 in an unpaired Student’s *t*-test. ns, not significant.

**Fig. 6.**
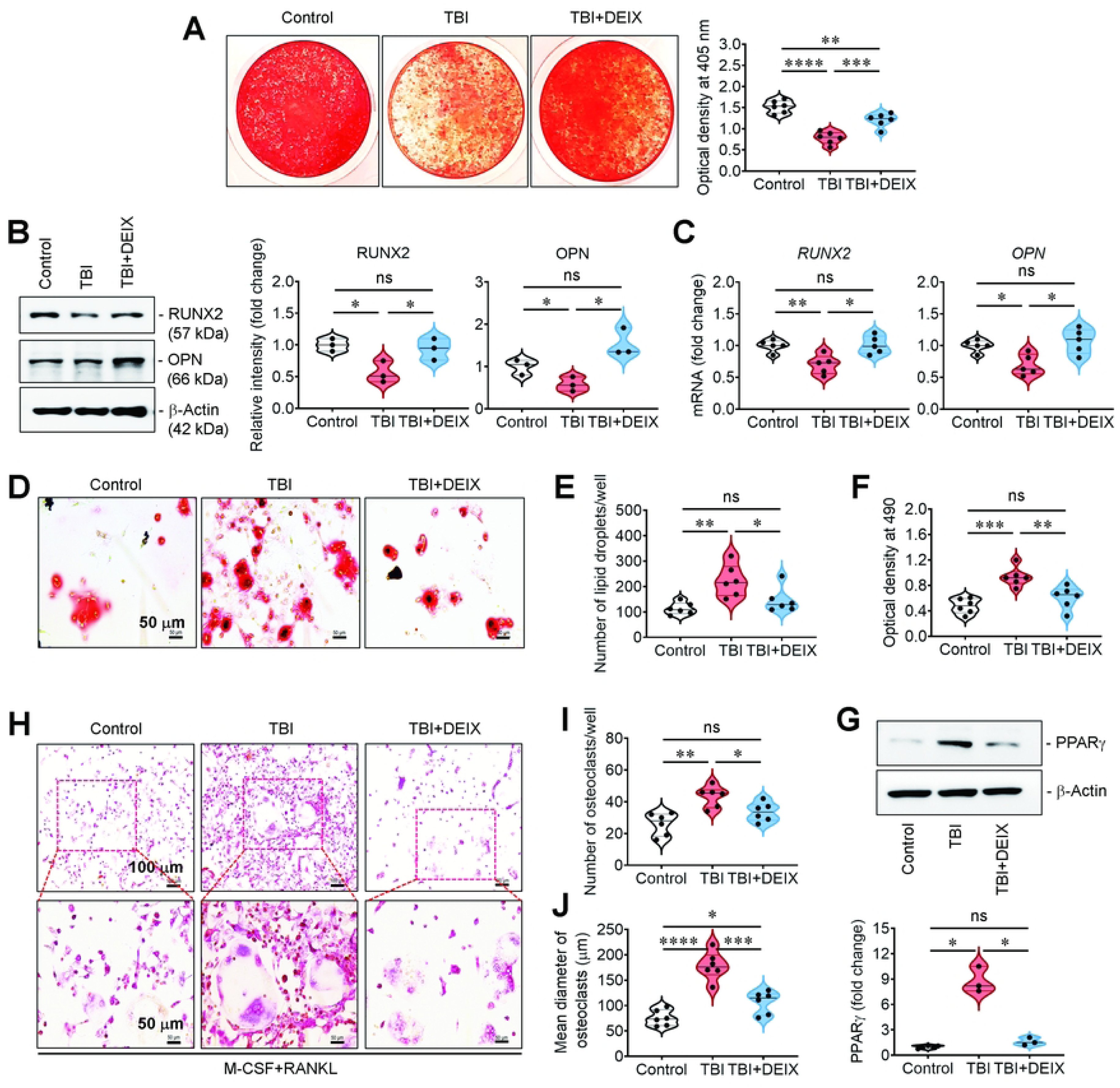
DEIX reverses TBI-mediated functional impairment in BMSCs and BMMs in their ability to differentiate into osteoblasts, adipocytes, or osteoclasts. BMSCs and BMMs were isolated from the mouse groups five weeks post-TBI, and their differentiation potential was analyzed. (**A**) ARS staining images showing the mineralization of BMSCs derived from the mouse groups and the dye-specific optical density at 405 nm (*n* = 6). (**B**) Western blot images showing the levels of RUNX2 and OPN in the mouse group-derived BMSCs five weeks post-TBI, in which levels (fold changes) of RUNX2 and OPN were determined after normalizing their levels to those of β-actin (*n* = 3). (**C**) RUNX2 and OPN levels in mouse group-derived BMSCs five weeks post-TBI, normalized to GAPDH (*n* = 5). (**D**) Photographs showing the formation of ORO-positive adipocytes by the mouse group-derived BMSCs. Determination of the (**E**) number of lipid droplets and (**F**) optical density corresponding to the ORO-specific dye after 21 days of incubation (*n* = 6). (**G**) Expression level of PPARγ in the mouse group-derived BM cells 35 days post-TBI (*n* = 3). (**H**) TRAP staining images showing osteoclasts formed by BMMs isolated from the mouse groups five days post-incubation. (**I**) Number and (**J**) mean diameter of osteoclasts that formed after osteoclastic stimulation with M-CSF and RANKL (*n* = 6). \**p* < 0.05, \*\**p* < 0.01, \*\*\**p* < 0.001, and \*\*\*\**p* < 0.0001 in an unpaired Student’s *t*-test. ns, not significant.

### Identification and functional analyses of DEGs via RNA sequence profiling

To better understand the role of supplemental DEIX in TBI-mediated BM impairments, we performed RNA sequencing on the mouse group-derived BM cells. The responses of DEGs and their KEGG and GO functional annotations were analyzed using the GEO database (accession GSE301666; https://www.ncbi.nlm.nih.gov/geo/query/acc.cgi?acc=GSE301666). Fig. 7A exhibits the heatmap of the DEGs selected from all samples of the mouse groups in log_10_ (FPKM+1) units and at the levels of fold change ≥ 2 and *p* value ≤ 0.05. We compared the numbers of DEGs between the control vs TBI or TBI+DEIX or DEIX and TBI vs TBI+DEIX group (Fig. 7B). In the control group, 78, 443, and 153 genes were upregulated, whereas 404, 662, and 355 genes were downregulated compared with the TBI, TBI+DEIX, and DEIX groups, at *q* ≤ 0.01, respectively. The TBI group showed 167 upregulated and 216 downregulated DEGs compared with the TBI+DEIX group. We conducted functional enrichment and pathway analysis of significantly enriched modules using g: Profiler and identified GO terms, including molecular function (MF), cellular compartment (CC), and biological process (BP), as well as KEGG pathways. Fig. 7C represents the plot summaries of GO and KEGG analyses between the indicated mouse groups, in which the DEGs in each of the indicated pathways were distributed as different colors in variously sized circles in -log10 (adjusted *p* value). Compared with the control group, the TBI group-derived BM cells showed a greater distribution of DEGs than the TBI+DEIX-derived cells. DEG numbers shown in the KEGG of the TBI group-derived BM cells were apparently reduced in the TBI+DEIX group-derived cells. Specifically, the DEIX group also identified a few DEGs associated with GO terms and KEGG pathways, compared with the control group. To better understand the effect of DEIX on the gene expression in TBI-exposed mice, the enriched biological terms from the DEGs between the control, TBI, and/or TBI+DEIX groups were divided into the top 10 terms in GO-BP, GO-CC, GO-MF, and KEGG pathways by singular enrichment analysis (SEA). The top 10 SEA terms of GO-BP (-10 log adjusted *p* value) were shown in Fig. 8A, in which the top one SEA in the control vs TBI groups was immune system process followed by the responses to other organism, external biotic stimulus, and biotic stimulus, etc. The top 10 SEA terms in GO-BP for the control and TBI+DEIX group-derived DEGs revealed the regulation of multicellular organismal process as the top one SEA term, followed by cell differentiation, cellular developmental process, and immune system process, etc. As in the GO-BP, the top DEG in the TBI vs TBI+DEIX group was an immune system process. However, the -10 log adjusted *p* value on the top 10 SEA terms derived from the control vs TBI group was apparently higher than that from the control vs TBI+DEIX group, indicating a restoration of TBI-derived DEGs by oral supplementation with DEIX. GO enrichment analysis for DEGs in the CC term revealed the external side of the plasma membrane, cell periphery, and membrane as the top SEA in the groups control vs TBI, control vs TBI+DEIX, and TBI vs TBI+DEIX, respectively (Fig. S1A). Similar to the GO-BP term, the -10 log adjusted *p* value of the GO-CC term for the control vs TBI group-derived DEGs was visibly lower compared with the value for the control vs TBI+DEIX group, or the TBI vs TBI+DEIX group-derived DEGs. However, the GO-MF analysis of DEGs showed protein binding as the top SEA term across all comparisons between the control, TBI, and TBI+DEIX groups, with no marked change in the -10 log adjusted *p* value (Fig. S1B). Alternatively, Fig. 8B shows the top 10 SEA terms of the KEGG pathway on the DEGs derived from the mouse groups. The DEGs between the control and TBI groups showed hematopoietic cell lineage as the top SEA, followed by terms associated with systemic and cellular immunological responses. Similarly, the SEA terms of the KEGG pathways from the DEGs of the control vs. TBI+DEIX group, or the TBI vs. TBI+DEIX group, revealed hematopoietic cell lineage as the top SEA. Unlike the top 10 SEA derived from the TBI group, the TBI+DEIX group showed two terms: hematopoietic cell lineage and primary immunodeficiency. To clarify the effect of supplemental DEIX on the TBI-mediated gene expression alterations, we screened 20 DEGs that showed 2-fold higher or lower expression in the TBI group compared with the control DEIX group. Fig. 8C shows the heatmap of the mean expression values for the selected DEGs across all samples with *p*-values < 0.05. The heatmap with a different color key indicated that a few DEGs are increased or decreased by TBI, whereas these changes are restored by DEIX administration. To further clarify the roles of supplemental DEIX on the BM-residing cells in TBI-exposed mice, we analyzed statistical differences of the DEGs in relation to TBI exposure and/or DEIX administration (Fig. 8D). Compared with the control group, the TBI group revealed significantly greater expression of *Apol11b*, *Col1a1*, *Col1a2, Ctsk*, *Fmo1*, *Sqle*, and *Tjp1* and lower expression in *Ak4*, *Bcl2a1a*, *Cd274*, *Ccl3*, *Ccl5*, *Cd3g*, *Cd8a*, *Col14a1*, *Csf1*, *IL4*, *Pdgfrb*, *Plcd3*, and *Prkcq*. No genes were differentially expressed between the control and TBI+DEIX groups, or between the control and DEIX groups, at significant levels. Taken together, these results indicate that supplemental DEIX restores or reverses the expression of genes upregulated or downregulated by TBI in the BM-retained cells.

**Fig. 7.**
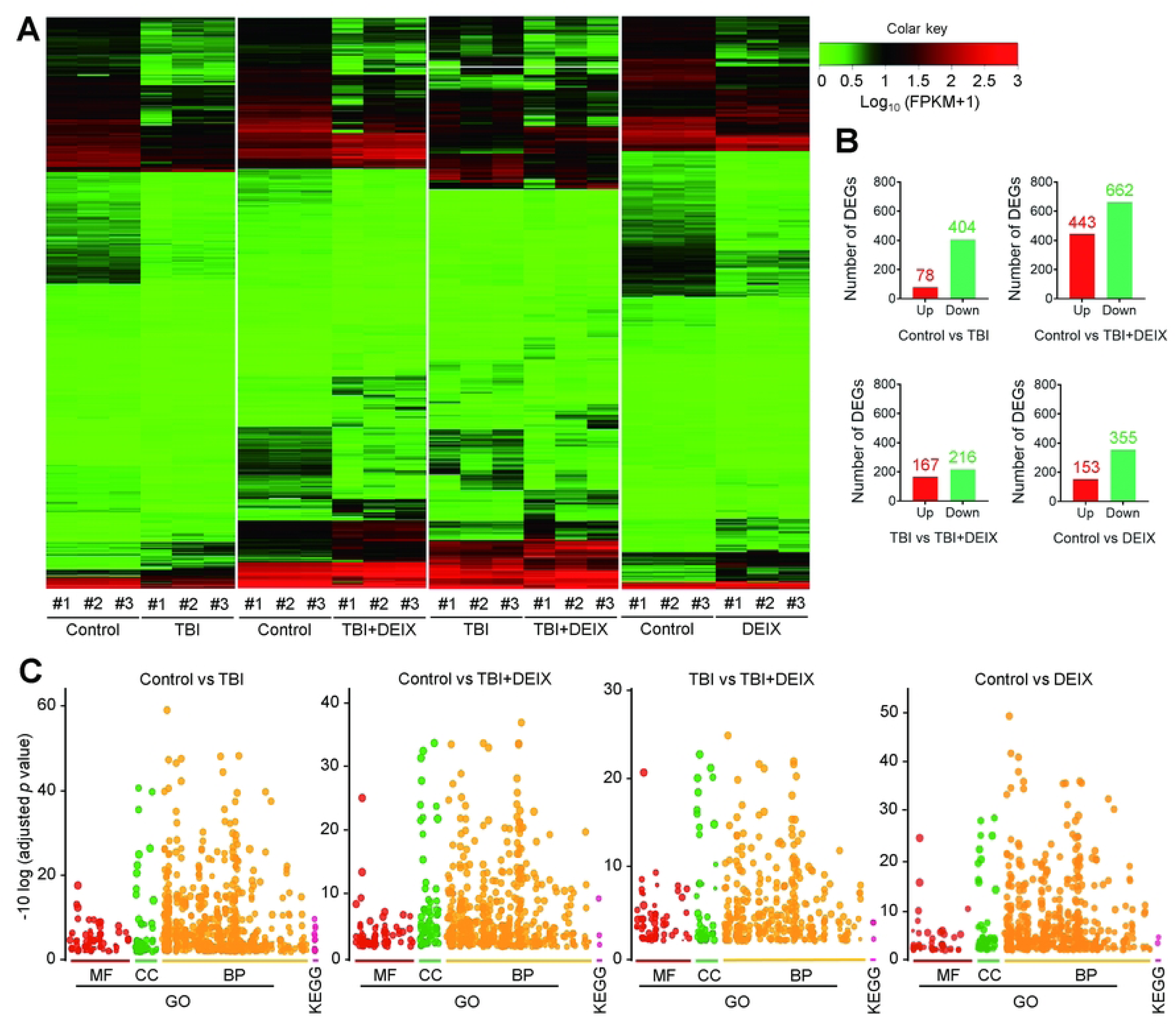
Identification and characterization of DEGs in whole BM cells of the mouse groups by RNA sequence profiling. (**A**) Cluster heatmap of representative DEGs in the mouse group-derived BM cells at the levels of fold change ≥ 2 and *p* value ≤ 0.05. (**B**) The graphs show the total numbers of DEGs from the indicated groups with *p*-values ≤ 0.05. (**C**) The plot summary images show the results of functional enrichment and pathway analysis of significantly enriched modules using g: Profiler, in which DEGs corresponding to GO-MF, GO-CC, and GO-BP, along with KEGG pathways, are represented by different colors and varying circle sizes, with -10 log-adjusted *p*-values.

**Fig. 8.**
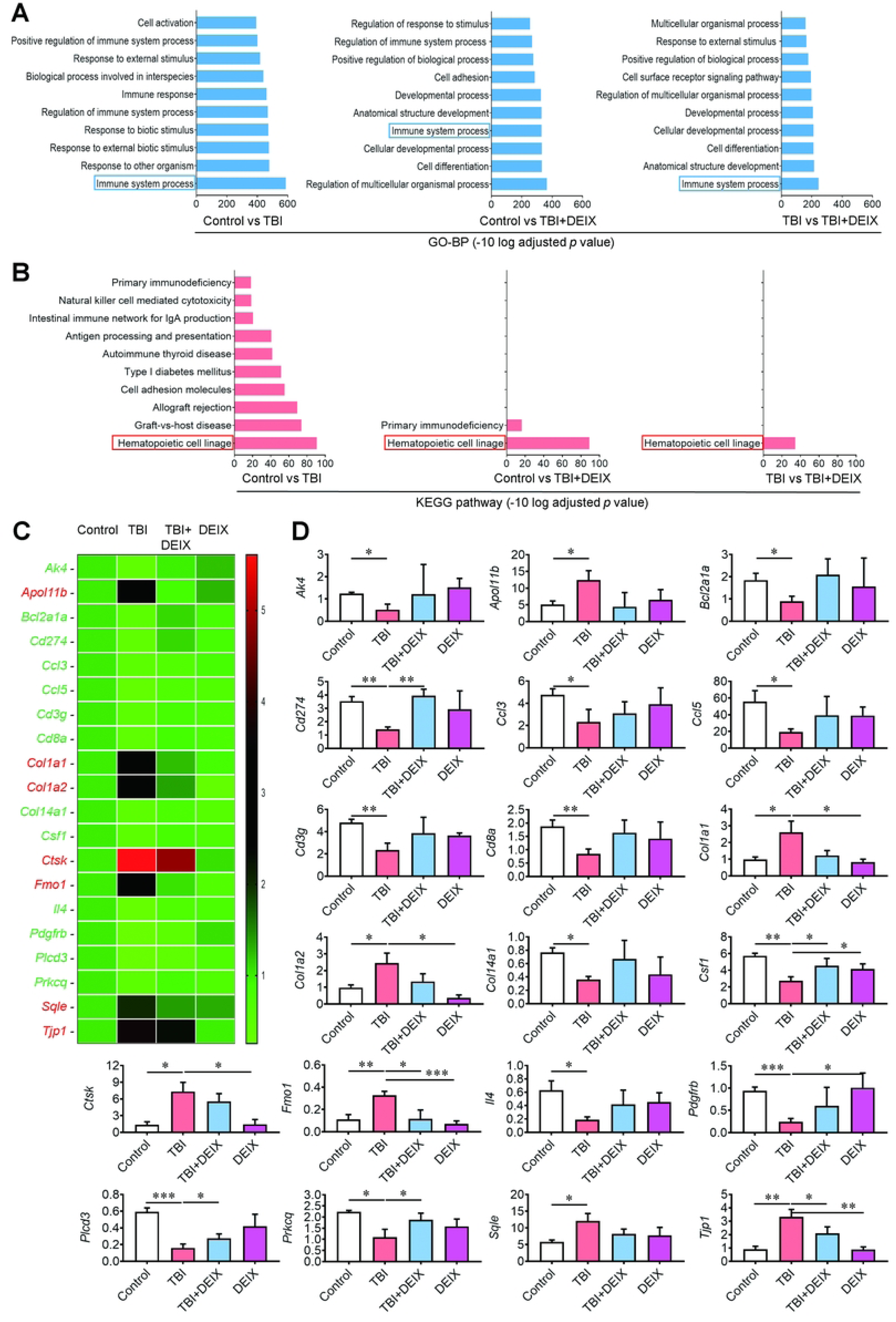
DEIX restores the TBI-induced expression of genes associated with various cellular events in BM-retained cells. Graphs exhibit the top 10 SEA terms of the indicated group-derived DEGs in (**A**) GO-BP and (**B**) KEGG pathways. The SEA terms marked in a blue or red box indicate the terms found in the control vs TBI+DEIX group, TBI vs TBI+DEIX, or both. The Y-axis displays the GO-BP and KEGG categories, each with up to 10 terms, ordered by -10 log-adjusted *p*-value. (**C**) The heatmap shows the comparative expression (fold change) of 20 selected genes in BM cells from the mouse groups in -10 log (FPKM+1) units. (**D**) The graphs show significant differences in DEGs between the indicated mouse group-derived BM cells (*p* < 0.05; *n* = 3). \**p* < 0.05, \*\**p* < 0.01, and \*\*\**p* < 0.001 in an unpaired Student’s *t*-test.

### The functional analyses of DEGs to determine the nature of DEIX on the gene expression in BM-retained cells

Based on RNA sequencing results, we further investigated the effects of DEIX treatment alone on gene expression in the mouse group-isolated BM cells. Among the top 10 SEAs derived from the control vs DEIX at -10 log-adjusted *p* value, the top terms for GO-BP (Fig. 9A), GO-CC (Fig. 9B), and GO-MF (Fig. 9C) were immune system process, side of membrane, and protein binding, respectively. Unlike GO terms, the KEGG pathway database identified two significantly enriched terms: primary immunodeficiency and hematopoietic cell lineage (Fig. 9D). From the DEIX-mediated DEGs, we selected 18 DEGs that showed significant differences from those of untreated controls at the level of *p* < 0.05. Fig. 9E shows the heatmap of the mean expression values for the selected DEGs across all samples, in which the expression levels of 12 up- and 6 down-regulated genes were compared in relation to the presence and absence of TBI exposure and/or DEIX administration. The results from a parametric Student *t*-test showed significantly higher expression of *Ddit4*, *Kcnj2*, *Lifr*, *Lrp1*, *Map3k6*, *Ptgfrn*, *Rufy4*, *Sla*, *Tgfbi*, *Tlr13*, *Tlr8*, and *Vldlr* in the DEIX group compared with the control, TBI, and/or TBI+DEIX groups (Fig. 9F). In contrast, the DEIX group showed significantly lower *Alpl*, *Cxcl12*, *Fabp4*, *Hpn*, *Ibsp*, and *Sphk1* levels compared with those in the control, TBI, and/or TBI+DEIX groups. These results indicate that supplemental DEIX itself affects the expression of genes closely associated with proliferation, metabolism, chemotaxis, differentiation, inflammation, and migration of BM-retained mesenchymal and hematopoietic lineage cells.

**Fig. 9.**
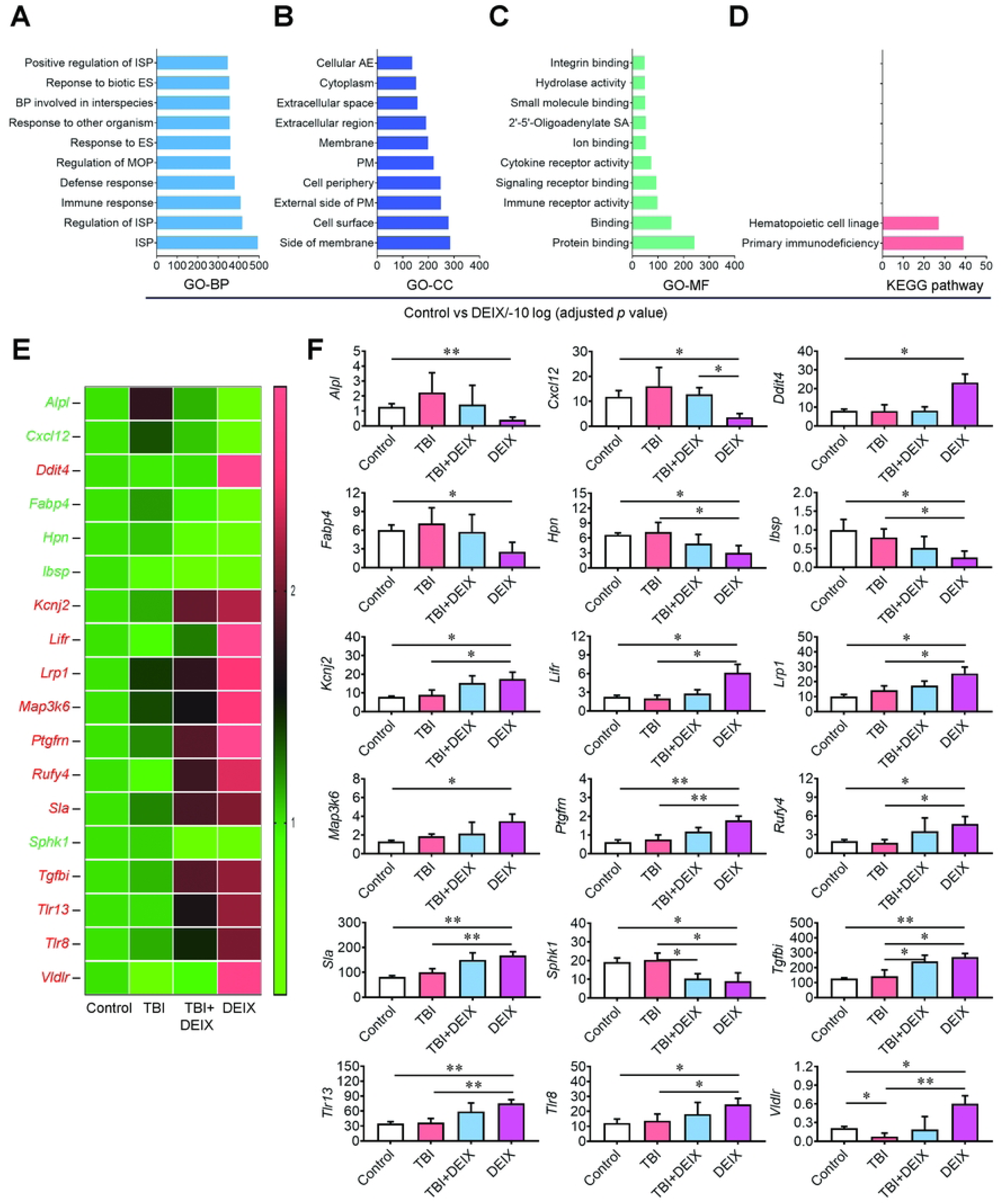
DEIX treatment alone affects the expression of genes associated with the fate, proliferation, survival, metabolism, and differentiation in BM cells. Significantly enriched top 10 terms of (**A**) GO-BP, (**B**) GO-CC, (**C**) GO-MF, and (**D**) KEGG pathway in the DEGs derived between the control and DEIX groups are shown. (**E**) The heatmap shows the expression levels of DEIX-specific DEGs relative to the control group, with 12 upregulated DEGs (red) and 6 downregulated DEGs (green). (**F**) The graphs show the significant differences of the selected DEGs between the indicated mouse group-derived BM cells at *p* < 0.05 (*n* = 3). \**p* < 0.05 and \*\**p* < 0.01 in an unpaired Student’s *t*-test. ISP, Immune system process; MOP, multicellular organismal process; ES, External stimulus; BP, Biological process; PM, plasma membrane; AE, anatomical entity; SA, synthetase activity.

## Discussion

TBI is a crucial treatment for cancer and BM transplantation patients, whereas it also causes severe damage to intact soft and hard tissues, depending on the doses and times applied [24, 25]. TBI-mediated damage includes decreases in bone mass and quality, excessive accumulation of intracellular ROS, long-term residual injury of BM, senescence and functional loss of BM-retained stem cells, and abnormal hematopoietic development [5, 26–28]. These TBI-mediated defects are closely associated with impairments in the BM microenvironment that provides niches for the retention and self-renewal of HSCs and MSCs [15, 27]. Defects in the activation of inflammation- and osteoclastogenesis-associated molecules, along with the downregulation of osteogenic molecules, are also associated with TBI-induced BM disorders [27, 29]. TBI-induced disorders in BM and BM-retained stem cells are similar to the hallmarks of age-related degenerative complications [30]. Consequently, our results strongly support the notion that TBI induces oxidative and senescence-associated complications in BM-residing stem cells and impairs the BM microenvironment.

Researchers have been developing bioactive materials to prevent or reverse TBI-mediated local and systemic damage without side effects. In this regard, we have investigated the radioprotective efficacies of naturally occurring phenolic antioxidants. For example, phenolic acids such as caffeic acid, ferulic acid, and coumaric acid protected against TBI-mediated mortality in an animal model by recovering TBI-induced oxidative injuries to BM and BM-retained cells [14, 27, 29]. A carotenoid compound, astaxanthin, protected against hyperglycemia-induced degenerative damage in the BM microenvironment and BM-retained stem cells [21]. Studies have also shown DEIX’s potency as a ROS scavenger and its anticancer and anti-inflammatory effects as a therapeutic antioxidant [18, 32–34]. Similarly, our previous findings showed that the addition of DEIX directly inhibits ROS accumulation and inflammatory responses in hydrogen peroxide-exposed BM cells, and oral DEIX supplementation may protect mice from TBI-mediated damage to growth, organs, and survival [19]. Accordingly, previous reports, along with our recent findings, support the notion that TBI-induced damage is accompanied by oxidative and inflammatory complications in BM and BM-residing cells, and that antioxidant compounds can mitigate these damages. Our current findings also demonstrate that DEIX supplementation protects against TBI-induced damage by ameliorating oxidative and inflammatory BM microenvironmental complications and by restoring the functionality of BM-retained stem cells.

Alternatively, BM adipocytes account for approximately 10% of the total body fat in healthy adults and play essential roles in energy storage, endocrine function, and bone metabolism [35]. Chemotherapy and irradiation can cause an abnormal infiltration and accumulation of adipocytes in the BM [9]. Such abnormal adipogenic activation in the BM could be associated with delayed HSC engraftment and hematopoietic defects [36]. TBI-mediated damage to MSC multipotency leads to preferential adipogenic differentiation, along with decreased osteogenesis and increased osteoclastogenesis in the BM [9]. TBI can arrest the cell cycle progression of osteoblastic progenitor cells and reduce the production of bone components [37]. Prolonged and persistent ROS accumulation in BM cells contributes to the induction of osteoclastogenesis, skeletal aging, and degenerative bone diseases [38]. Consequently, our ex vivo findings indicate that TBI-mediated BM injuries result from dysregulated differentiation or function of MSC-derived progenitor cells, and that these injuries are recoverable with long-term supplementation with DEIX.

To better understand the mechanisms underlying TBI-induced BM impairment and its inhibition by supplemental DEIX, we performed RNA sequencing on the mouse group-derived BM cells. Analyzing the DEGs with GO functional analysis and KEGG pathways indicates that TBI-mediated changes primarily affect cellular and systemic immune responses, as well as hematopoietic development. The comparative study of the 20 selected DEGs further supports the notion of TBI-mediated immunological disorders and their restoration by supplemental DEIX. Of the 20 chosen DEGs, the *Ak4* gene encodes an enzyme that regulates cellular nucleotide homeostasis [39], while *Apol11b* encodes a protein component of lipid metabolism-involved lipoproteins [40]. *Bcl2a1a*encodes an anti-apoptotic protein and plays a crucial role in hematopoiesis [41]. The *Ccl3* and *Ccl5* genes encode chemokines that play central roles in the immune system by recruiting or modulating immune cells [42]. The *Cd3* encodes a critical component of the T-cell receptor/CD3 complex that initiates T-cell activation [43], and the *Cd8* encodes a key element required for the activation of cytotoxic T lymphocytes [44]. The genes *Col1a1* and *Col1a2* encode α-1 and α-2 type I collagens, which are found in most connective tissues, whereas *Col14a1* plays a role in collagen binding and cell-cell adhesion [45, 46]. While the *Csf1*encodes a secreted cytokine, M-CSF (also known as CSF1) that stimulates hematopoietic differentiation [47], *Ctsk* encodes cathepsin K that mediates bone remodeling and resorption [48]. The *Il4* gene encodes interleukin 4 (IL4) that induces differentiation of naïve helper T cells and contributes to the function of T and B cells [49]. The *Pdgfrb* gene encodes platelet-derived growth factor receptor β, which plays an essential role in vascular development [50]. In contrast, the *Plcd3* gene encodes an enzyme that produces intracellular second messengers that are required for various biological processes [51]. The *Prkcq* gene encodes protein kinase C, and is involved in neuronal function and immune responses [52]. The *Sqle* gene encodes squalene epoxidase, which is responsible for cholesterol biosynthesis [53], whereas *Tjp1* encodes tight junction protein 1, which maintains tight junction integrity and modulates cellular processes, such as development, migration, and signal transduction [54]. Taken together, our results suggest that TBI-mediated BM complications and their improvement with supplemental DEIX may involve the restoration of gene expression for enzymes, chemokines, and cytokines essential for cellular and systemic immune modulation and various biological processes.

The RNA sequencing profiling on the mouse group-isolated BM cells also indicates that DEIX supplementation alone may upregulate the genes involved in MSC fate and erythroid differentiation (*Ddit4*) [55, 56], cell development and osteoblast function (*Kcnj2*) [57], stem cell maintenance and bone remodeling (*Lifr*) [58], bone resorption and formation (*Lrp1*) [59], survival and stress response (*Map3k6*) [60], HSC maintenance (*Ptgfrn*) [61], bone resorption and autophagy regulation (*Rufy4*) [62], BM microenvironment controlling (*Sla*) [63], cell survival, adhesion, and differentiation balance (*Tgfbi*) [64], macrophage activation (*Tlr13*) [65], immune sensing and inflammatory signaling (*Tlr8*) [66], and osteoclast regulation (*Vldlr*) [67]. In contrast, the sequencing analysis postulates that the genes encoding the proteins involved in bone metabolism and stem cell regulation (*Alpl*) [68], HSC retention and homing (*Cxcl12*) [69], bone resorption promotion (*Fabp4*) [70], cancer cell metastasis facilitation (*Hpn*) [71], bone mineralization (*Ibsp*; also known as bone sialoprotein) [72], and osteoblast coupling (*Sphk1*) [73], can be downregulated in BM cells of the DEIX-only-supplied mice. Collectively, these findings support the notion that DEIX treatment alone exhibits beneficial roles on the expression of genes associated with the fate of BM cells and the maintenance of the BM microenvironment. However, additional experiments using multiple numbers of animal models might be needed to further clarify the protective efficacy and the exact mechanisms of supplemental DEIX in BM and BM cells of TBI-exposed mice, as well as to evaluate its clinical usefulness for patients who require TBI.

## Conclusions

In summary, supplemental DEIX protects mice from TBI-mediated BM injury and the senescence of BM-retained mesenchymal lineage cells. That protection is orchestrated by inhibiting ROS accumulation and adipocytic activation in BM, as well as by restoring the imbalance between osteoblast and osteoclast activity and antioxidant defense system (Fig. 10). Collectively, our results indicate that DEIX has a clinical potential to prevent TBI-mediated oxidative and inflammatory disorders in BM and BM-retained cells and enhance the therapeutic efficacy of BM transplantation.

**Fig. 10.**
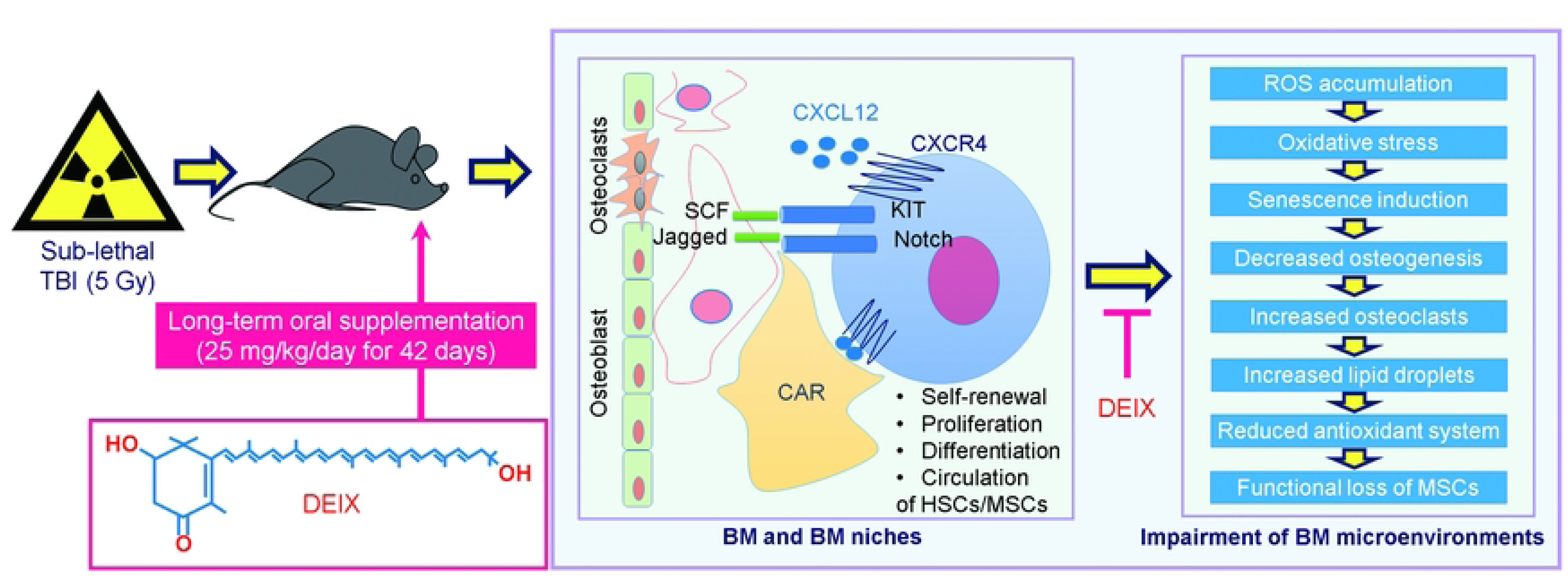
**Schematic illustrations showing the radioprotective mechanisms of DEIX in TBI-exposed mice.**

## Supporting information

**Fig. S1** Significantly enriched top 10 terms of (**A**) GO-CC and (**B**) GO-MF in the DEGs between the indicated mouse group-derived BM cells.

## Author Contributions

Conceptualization: Shankar Rijal, Kihyun Kim, Govinda Bhattarai, Junhyeok Kim, Bitna Kim, Young-Mi Jeon, Sung-Ho Kook, Jeong-Chae Lee.

Data curation: Shankar Rijal, Kihyun Kim, Govinda Bhattarai, Sung-Ho Kook, Sung-Ho Kook, Jeong-Chae Lee.

Formal analysis: Shankar Rijal, Kihyun Kim, Govinda Bhattarai, Junhyeok Kim, Bitna Kim, Young-Mi Jeon, Sung-Ho Kook, Jeong-Chae Lee.

Funding acquisition: Govinda Bhattarai, Young-Mi Jeon, Sung-Ho Kook, Jeong-Chae Lee. Investigation: Shankar Rijal, Kihyun Kim, Govinda Bhattarai, Junhyeok Kim, Bitna Kim. Methodology: Shankar Rijal, Kihyun Kim, Govinda Bhattarai, Junhyeok Kim, Bitna Kim, Sung-Ho Kook, Jeong-Chae Lee.

Validation: Shankar Rijal, Kihyun Kim, Junhyeok Kim.

Project administration: Young-Mi Jeon, Sung-Ho Kook, Jeong-Chae Lee. Supervision: Sung-Ho Kook, Jeong-Chae Lee.

Visualization: Shankar Rijal, Kihyun Kim, Govinda Bhattarai.

Writing – original draft: Shankar Rijal, Kihyun Kim, Govinda Bhattarai. Writing – review and editing: Sung-Ho Kook, Jeong-Chae Lee.

## Acknowledgments

This work was supported by the National Research Foundation (NRF) of Korea, funded by the Ministry of Science, Information and Communications Technology and Future Planning [RS-2023-00277774, RS-2024-00338143, RS-2026-25473404, and RS-2026-25474928].

## Data availability

The mRNA sequence data supporting the findings of this study have been deposited in the GEO database under accession GSE301666 (https://www.ncbi.nlm.nih.gov/geo/query/acc.cgi?acc=GSE301666).

## References

1. Kasbekar M, Mitchell CA, Proven MA, Passegué E. Hematopoietic stem cells through the ages: A lifetime of adaptation to organismal demands. Cell Stem Cell. 2023; 30: 1403–1420. doi:10.1016/j.stem.2023.09.013

2. Melin N, Yarahmadov T, Sanchez-Taltavull D, Birrer FE, Brodie TM, Petit B, et al. A new mouse model of radiation-induced liver disease reveals mitochondrial dysfunction as an underlying fibrotic stimulus. JHEP Rep. 2022; 4: 100508. doi:10.1016/j.jhepr.2022.100508

3. Shao L, Wang Y, Chang J, Luo Y, Meng A, Zhou D. Hematopoietic stem cell senescence and cancer therapy-induced long-term bone marrow injury. Transl Cancer Res. 2013; 2: 397–411. doi:10.3978/j.issn.2218-676X.2013.07.03

4. Bai J, Wang Y, Wang J, Zhai J, He F, Zhu G. Irradiation-induced senescence of bone marrow mesenchymal stem cells aggravates osteogenic differentiation dysfunction via paracrine signaling. Am J Physiol Cell Physiol. 2020; 318: C1005–C1017. doi:10.1152/ajpcell.00520.2019

5. Shao L, Feng W, Li H, Gardner D, Luo Y, Wang Y, et al. Total body irradiation causes long-term mouse BM injury via induction of HSC premature senescence in an Ink4a- and Arf-independent manner. Blood. 2014; 123: 3105–3115. doi:10.1182/blood-2013-07-515619

6. Wang Y, Schulte BA, LaRue AC, Ogawa M, Zhou D. Total body irradiation selectively induces murine hematopoietic stem cell senescence. Blood. 2006; 107: 358–366. doi:10.1182/blood-2005-04-1418

7. Wang Y, Liu L, Pazhanisamy SK, Li H, Meng A, Zhou D. Total body irradiation causes residual bone marrow injury by induction of persistent oxidative stress in murine hematopoietic stem cells. Free Radic Biol Med. 2010; 48: 348–356. doi:10.1016/j.freeradbiomed.2009.11.005

8. Cao X, Wu X, Frassica D, Yu B, Pang L, Xian L, et al. Irradiation induces bone injury by damaging bone marrow microenvironment for stem cells. Proc Natl Acad Sci USA. 2011; 108: 1609–1614. doi:10.1073/pnas.1015350108

9. Costa S, Reagan MR. Therapeutic Irradiation: Consequences for Bone and Bone Marrow Adipose Tissue. Front Endocrinol (Lausanne). 2019; 10: 587. doi:10.3389/fendo.2019.00587

10. Das U, Manna K, Sinha M, Datta S, Das DK, Chakraborty A, et al. Role of ferulic acid in the amelioration of ionizing radiation induced inflammation: a murine model. PLoS One. 2014; 9: e97599. doi:10.1371/journal.pone.0097599

11. Li D, Rui YX, Guo SD, Luan F, Liu R, Zeng N. Ferulic acid: A review of its pharmacology, pharmacokinetics and derivatives. Life Sci. 2021; 284: 119921. doi:10.1016/j.lfs.2021.119921

12. Piazzon A, Vrhovsek U, Masuero D, Mattivi F, Mandoj F, Nardini M. Antioxidant activity of phenolic acids and their metabolites: synthesis and antioxidant properties of the sulfate derivatives of ferulic and caffeic acids and of the acyl glucuronide of ferulic acid. J Agric Food Chem. 2012; 60: 12312–12323. doi:10.1021/jf304076z

13. Rudrapal M, Khairnar SJ, Khan J, Dukhyil AB, Ansari MA, Alomary MN, et al. Dietary Polyphenols and Their Role in Oxidative Stress-Induced Human Diseases: Insights Into Protective Effects, Antioxidant Potentials and Mechanism(s) of Action. Front Pharmacol. 2022; 13: 806470. doi:10.3389/fphar.2022.806470

14. Wagle S, Sim HJ, Bhattarai G, Choi KC, Kook SH, Lee JC, Jeon YM. Supplemental Ferulic Acid Inhibits Total Body Irradiation-Mediated Bone Marrow Damage, Bone Mass Loss, Stem Cell Senescence, and Hematopoietic Defect in Mice by Enhancing Antioxidant Defense Systems. Antioxidants (Basel). 2021; 10: 1209. doi:10.3390/antiox10081209

15. Liang JW, Li PL, Wang Q, Liao S, Hu W, Zhao ZD, et al. Ferulic acid promotes bone defect repair after radiation by maintaining the stemness of skeletal stem cells. Stem Cells Transl Med. 2021; 10: 1217–1231. doi:10.1002/sctm.20-0536

16. Wang C, Zhao S, Shao X, Park JB, Jeong SH, Park HJ, et al. Challenges and tackles in metabolic engineering for microbial production of carotenoids. Microb Cell Fact. 2019; 18: 55. doi:10.1186/s12934-019-1105-1

17. Tian B, Xu Z, Sun Z, Lin J, Hua Y. Evaluation of the antioxidant effects of carotenoids from *Deinococcus radiodurans* through targeted mutagenesis, chemiluminescence, and DNA damage analyses. Biochim Biophys Acta. 2007; 1770: 902–911. doi:10.1016/j.bbagen.2007.01.016

18. Tian B, Hua Y. Carotenoid biosynthesis in extremophilic *Deinococcus-Thermus* bacteria. Trends Microbiol. 2010; 18: 512–520. doi:10.1016/j.tim.2010.07.007

19. Bhattarai G, Kook SH, Shrestha SK, Park JH, Rijal S, Tae G, et al. Deinoxanthin Recovers H_2_O_2_-Stimulated Oxidative Complications of Bone Marrow-Derived Cells and Protects Mice from Irradiation-Mediated Impairments. Antioxidants (Basel). 2025; 14: 1180. doi:10.3390/antiox14101180

20. Bhattarai G, An YH, Shrestha SK, Rijal S, Park SM, Kook SH, et al. Therapeutic potency and the related mechanism of deinoxanthin in experimental animal and cell models of periodontitis. Sci Rep. 2026; 16: 5735. doi:10.1038/s41598-026-36514-1

21. Bhattarai G, So HS, Kieu TTT, Kook SH, Lee JC, Jeon YM. Astaxanthin Inhibits Diabetes-Triggered Periodontal Destruction, Ameliorates Oxidative Complications in STZ-Injected Mice, and Recovers Nrf2-Dependent Antioxidant System. Nutrients. 2021; 13: 3575. doi:10.3390/nu13103575

22. Kiel MJ, Yilmaz OH, Iwashita T, Yilmaz OH, Terhorst C, Morrison SJ. SLAM family receptors distinguish hematopoietic stem and progenitor cells and reveal endothelial niches for stem cells. Cell. 2005; 121: 1109–1121. doi:10.1016/j.cell.2005.05.026

23. Sim HJ, So HS, Poudel SB, Bhattarai G, Cho ES, Lee JC, Kook SH. Osteoblastic Wls Ablation Protects Mice from Total Body Irradiation-Induced Impairments in Hematopoiesis and Bone Marrow Microenvironment. Aging Dis. 2023; 14: 919–936. doi:10.14336/AD.2022.1026

24. King AD, Griffith JF, Abrigo JM, Leung SF, Yau FK, Tse GM, Ahuja AT. Osteoradionecrosis of the upper cervical spine: MR imaging following radiotherapy for nasopharyngeal carcinoma. Eur J Radiol. 2010; 73: 629–635. doi:10.1016/j.ejrad.2008.12.016

25. Schultze-Mosgau S, Lehner B, Rödel F, Wehrhan F, Amann K, Kopp J, et al. Expression of bone morphogenic protein 2/4, transforming growth factor-beta1, and bone matrix protein expression in healing area between vascular tibia grafts and irradiated bone-experimental model of osteonecrosis. Int J Radiat Oncol Biol Phys. 2005; 61: 1189–1196. doi:10.1016/j.ijrobp.2004.12.008

26. Lo WJ, Lin CL, Chang YC, Bai LY, Lin CY, Liang JA, et al. Total body irradiation tremendously impair the proliferation, differentiation and chromosomal integrity of bone marrow-derived mesenchymal stromal stem cells. Ann Hematol. 2018; 97: 697–707. doi:10.1007/s00277-018-3231-y

27. Sim HJ, Bhattarai G, Lee J, Lee JC, Kook SH. The Long-lasting Radioprotective Effect of Caffeic Acid in Mice Exposed to Total Body Irradiation by Modulating Reactive Oxygen Species Generation and Hematopoietic Stem Cell Senescence-Accompanied Long-term Residual Bone Marrow Injury. Aging Dis. 2019; 10: 1320–1327. doi:10.14336/AD.2019.0208

28. Wang Y, Zhu G, Wang J, Chen J. Irradiation alters the differentiation potential of bone marrow mesenchymal stem cells. Mol Med Rep. 2016; 13: 213–223. doi:10.3892/mmr.2015.4539

29. Koc M, Taysi S, Buyukokuroglu ME, Bakan N. Melatonin protects rat liver against irradiation-induced oxidative injury. J Radiat Res. 2003; 44: 211–215. doi:10.1269/jrr.44.211

30. Zupan J, Strazar K, Kocijan R, Nau T, Grillari J, Marolt Presen D. Age-related alterations and senescence of mesenchymal stromal cells: Implications for regenerative treatments of bones and joints. Mech Ageing Dev. 2021; 198: 111539. doi:10.1016/j.mad.2021.111539

31. Kook SH, Cheon SR, Kim JH, Choi KC, Kim MK, Lee JC. Dietary hydroxycinnamates prevent oxidative damages to liver, spleen, and bone marrow cells in irradiation-exposed mice. Food Sci Biotechnol. 2017; 26: 279–285. doi:10.1007/s10068-017-0037-y

32. Choi YJ, Hur JM, Lim S, Jo M, Kim DH, Choi JI. Induction of apoptosis by deinoxanthin in human cancer cells. Anticancer Res. 2014; 34: 1829–1835. PMID: 24692716

33. Maqbool I, Sudharsan M, Kanimozhi G, Alrashood ST, Khan HA, Prasad NR. Crude Cell-Free Extract From *Deinococcus radiodurans* Exhibit Anticancer Activity by Inducing Apoptosis in Triple-Negative Breast Cancer Cells. Front Cell Dev Biol. 2020; 8: 707. doi:10.3389/fcell.2020.00707

34. Yu S, Kim S, Kim J, Kim JW, Kim SY, Yeom B, et al. Highly Water-Dispersed and Stable Deinoxanthin Nanocapsule for Effective Antioxidant and Anti-Inflammatory Activity. Int J Nanomedicine. 2023; 18: 4555–4565. doi:10.2147/IJN.S401808

35. Herrmann M. Marrow Fat-Secreted Factors as Biomarkers for Osteoporosis. Curr Osteoporos Rep. 2019; 17: 429–437. doi:10.1007/s11914-019-00550-w

36. Naveiras O, Nardi V, Wenzel PL, Hauschka PV, Fahey F, Daley GQ. Bone-marrow adipocytes as negative regulators of the haematopoietic microenvironment. Nature. 2009; 460: 259–263. doi:10.1038/nature08099

37. Johnson MB, Niknam-Bienia S, Soundararajan V, Pang B, Jung E, Gardner DJ, et al. Mesenchymal Stromal Cells Isolated from Irradiated Human Skin Have Diminished Capacity for Proliferation, Differentiation, Colony Formation, and Paracrine Stimulation. Stem Cells Transl Med. 2019; 8: 925–934. doi:10.1002/sctm.18-0112

38. Marques-Carvalho A, Kim HN, Almeida M. The role of reactive oxygen species in bone cell physiology and pathophysiology. Bone Rep. 2023; 19: 101664. doi:10.1016/j.bonr.2023.101664

39. Fujisawa K, Terai S, Takami T, Yamamoto N, Yamasaki T, Matsumoto T, et al. Modulation of anti-cancer drug sensitivity through the regulation of mitochondrial activity by adenylate kinase 4. J Exp Clin Cancer Res. 2016; 35: 48. doi:10.1186/s13046-016-0322-2

40. Raulin AC, Martens YA, Bu G. Lipoproteins in the Central Nervous System: From Biology to Pathobiology. Annu Rev Biochem. 2022; 91: 731–759. doi:10.1146/annurev-biochem-032620-104801

41. Métais JY, Winkler T, Geyer JT, Calado RT, Aplan PD, Eckhaus MA, Dunbar CE. BCL2A1a over-expression in murine hematopoietic stem and progenitor cells decreases apoptosis and results in hematopoietic transformation. PLoS One. 2012; 7: e48267. doi:10.1371/journal.pone.0048267

42. Ozga AJ, Chow MT, Luster AD. Chemokines and the immune response to cancer. Immunity. 2021; 54: 859–874. doi:10.1016/j.immuni.2021.01.012

43. Zheng J, Zhang Y, Cai Y, Han W, Chen W. An optimized non-T cell transfection system based on HEK293FT cells for CD3ζ phosphorylation and ubiquitination. J Immunol Methods. 2024; 528: 113664. doi:10.1016/j.jim.2024.113664

44. Sykulev Y. Factors contributing to the potency of CD8^+^ T cells. Trends Immunol. 2023; 44: 693–700. doi:10.1016/j.it.2023.07.005

45. Chen Y, Yang S, Lovisa S, Ambrose CG, McAndrews KM, Sugimoto H, Kalluri R. Type-I collagen produced by distinct fibroblast lineages reveals specific function during embryogenesis and Osteogenesis Imperfecta. Nat Commun. 2021; 12: 7199. doi:10.1038/s41467-021-27563-3

46. Skov V, Thomassen M, Kjaer L, Riley C, Larsen TS, Bjerrum OW, et al. Extracellular Matrix-Related Genes Are Deregulated in Peripheral Blood from Patients with Myelofibrosis and Related Neoplasms. Blood. 2018; 132: 5491. doi:10.1182/blood-2018-99-117122

47. Wang Q, Wang J, Xu K, Luo Z. Targeting the CSF1/CSF1R signaling pathway: an innovative strategy for ultrasound combined with macrophage exhaustion in pancreatic cancer therapy. Front Immunol. 2024; 15: 1481247. doi:10.3389/fimmu.2024.1481247

48. Wu N, Wang Y, Wang K, Zhong B, Liao Y, Liang J, Jiang N. Cathepsin K regulates the tumor growth and metastasis by IL-17/CTSK/EMT axis and mediates M2 macrophage polarization in castration-resistant prostate cancer. Cell Death Dis. 2022; 13: 813. doi:10.1038/s41419-022-05215-8

49. Ul-Haq Z, Naz S, Mesaik MA. Interleukin-4 receptor signaling and its binding mechanism: A therapeutic insight from inhibitors tool box. Cytokine Growth Factor Rev. 2016; 32: 3–15. doi:10.1016/j.cytogfr.2016.04.002

50. Ando K, Shih YH, Ebarasi L, Grosse A, Portman D, Chiba A, et al. Conserved and context-dependent roles for pdgfrb signaling during zebrafish vascular mural cell development. Dev Biol. 2021; 479: 11–22. doi:10.1016/j.ydbio.2021.06.010

51. Liu W, Liu X, Wang L, Zhu B, Zhang C, Jia W, et al. PLCD3, a flotillin2-interacting protein, is involved in proliferation, migration and invasion of nasopharyngeal carcinoma cells. Oncol Rep. 2018; 39: 45–52. doi:10.3892/or.2017.6080

52. Saito N, Shirai Y. Protein kinase C gamma (PKC gamma): function of neuron specific isotype. J Biochem. 2002; 132: 683–687. doi:10.1093/oxfordjournals.jbchem.a003274

53. Zhang L, Cao Z, Hong Y, He H, Chen L, Yu Z, Gao Y. Squalene Epoxidase: Its Regulations and Links with Cancers. Int J Mol Sci. 2024; 25: 3874. doi:10.3390/ijms25073874

54. Zhang XD, Baladandayuthapani V, Lin H, Mulligan G, Li B, Esseltine DW, et al. Tight Junction Protein 1 Modulates Proteasome Capacity and Proteasome Inhibitor Sensitivity in Multiple Myeloma via EGFR/JAK1/STAT3 Signaling. Cancer Cell. 2016; 29: 639–652. doi:10.1016/j.ccell.2016.03.026

55. Cui Y, Zhou XY, Li XX, Yang YD, Yang CZ, Chen DW, et al. DDIT4 promotes erythroid differentiation and coordinates with SIPA1 to regulate erythroid proliferation in bone marrow of high altitude erythrocytosis. Life Sci. 2024; 359: 123212. doi:10.1016/j.lfs.2024.123212

56. Gharibi B, Ghuman M, Hughes FJ. DDIT4 regulates mesenchymal stem cell fate by mediating between HIF1α and mTOR signalling. Sci Rep. 2016; 6: 36889. doi:10.1038/srep36889

57. Shen M, Pan R, Lei S, Zhang L, Zhou C, Zeng Z, et al. KCNJ2/HIF1α positive-feedback loop promotes the metastasis of osteosarcoma. Cell Commun Signal. 2023; 21: 46. 10.1186/s12964-023-01064-w

58. Fierro FA, Nolta JA, Adamopoulos IE. Concise Review: Stem Cells in Osteoimmunology. Stem Cells. 2017; 35: 1461–1467. doi:10.1002/stem.2625

59. Bartelt A, Behler-Janbeck F, Beil FT, Koehne T, Müller B, Schmidt T, et al. Lrp1 in osteoblasts controls osteoclast activity and protects against osteoporosis by limiting PDGF-RANKL signaling. Bone Res. 2018; 6: 4. doi:10.1038/s41413-017-0006-3

60. Eto N, Miyagishi M, Inagi R, Fujita T, Nangaku M. Mitogen-activated protein 3 kinase 6 mediates angiogenic and tumorigenic effects via vascular endothelial growth factor expression. Am J Pathol. 2009; 174: 1553–1563. doi:10.2353/ajpath.2009.080190

61. Himburg HA, Harris JR, Ito T, Daher P, Russell JL, Quarmyne M, et al. Pleiotrophin regulates the retention and self-renewal of hematopoietic stem cells in the bone marrow vascular niche. Cell Rep. 2012; 2: 964–975. doi:10.1016/j.celrep.2012.09.002

62. Kim M, Park JH, Go M, Lee N, Seo J, Lee H, et al. RUFY4 deletion prevents pathological bone loss by blocking endo-lysosomal trafficking of osteoclasts. Bone Res. 2024; 12: 29. doi:10.1038/s41413-024-00326-8

63. Kazi JU, Kabir NN, Rönnstrand L. Role of SRC-like adaptor protein (SLAP) in immune and malignant cell signaling. Cell Mol Life Sci. 2015; 72: 2535–2544. doi:10.1007/s00018-015-1882-6

64. Klamer SE, Dorland YL, Kleijer M, Geerts D, Lento WE, van der Schoot CE, et al. TGFBI Expressed by Bone Marrow Niche Cells and Hematopoietic Stem and Progenitor Cells Regulates Hematopoiesis. Stem Cells Dev. 2018; 27: 1494–1506. doi:10.1089/scd.2018.0124

65. Signorino G, Mohammadi N, Patanè F, Buscetta M, Venza M, Venza I, et al. Role of Toll-like receptor 13 in innate immune recognition of group B streptococci. Infect Immun. 2014; 82: 5013–5022. doi:10.1128/IAI.02282-14

66. Cervantes JL, Weinerman B, Basole C, Salazar JC. TLR8: the forgotten relative revindicated. Cell Mol Immunol. 2012; 9: 434–438. doi:10.1038/cmi.2012.38

67. Huynh H, Wei W, Wan Y. mTOR Inhibition Subdues Milk Disorder Caused by Maternal VLDLR Loss. Cell Rep. 2017; 19: 2014–2025. doi:10.1016/j.celrep.2017.05.037

68. Liu W, Zhang L, Xuan K, Hu C, Liu S, Liao L, et al. *Alpl* prevents bone ageing sensitivity by specifically regulating senescence and differentiation in mesenchymal stem cells. Bone Res. 2018; 6: 27. doi:10.1038/s41413-018-0029-4

69. Yellowley C. CXCL12/CXCR4 signaling and other recruitment and homing pathways in fracture repair. Bonekey Rep. 2013; 2: 300. doi:10.1038/bonekey.2013.34

70. Pan C, Li S, Li C, Ko WC, Liu T, Liu H, et al. Fabkin Promoted Osteoclasts Mature and Bone Loss in OVX-Induced Osteoporosis Mice. FASEB J. 2025; 39: e71028. doi:10.1096/fj.202501946R

71. Klezovitch O, Chevillet J, Mirosevich J, Roberts RL, Matusik RJ, Vasioukhin V. Hepsin promotes prostate cancer progression and metastasis. Cancer Cell. 2004; 6: 185–195. 10.1016/j.ccr.2004.07.008

72. Kottmann V, Drees P, Gercek E, Ritz U. Bone sialoprotein: a multifunctional regulator of bone remodelling and tumour progression. Bone Res. 2026; 14: 11. doi:10.1038/s41413-025-00490-5

73. Ryu J, Kim HJ, Chang EJ, Huang H, Banno Y, Kim HH. Sphingosine 1-phosphate as a regulator of osteoclast differentiation and osteoclast-osteoblast coupling. EMBO J. 2006; 25: 5840–5851. doi:10.1038/sj.emboj.7601430

